# Stretch and flow at the gliovascular interface: high-fidelity modelling of astrocyte endfeet

**DOI:** 10.1101/2025.05.08.652799

**Authors:** Marius Causemann, Rune Enger, Marie E. Rognes

## Abstract

Astrocyte endfeet form a near-continuous sheath around the brain’s vasculature, defining the perivascular spaces (PVS) that are crucial for brain fluid flow and solute transport. Yet, their precise physiological role remains poorly understood. Using 3D electron microscopy data, we created a high-fidelity poroelastic computational model of an arteriole segment with surrounding endfeet and parenchyma to investigate tissue displacement and fluid flow within the PVS, endfeet, and extracellular space (ECS) in response to blood vessel pulsations. Our model predicts that arteriole dilations compress the PVS while expanding the overall endfoot sheath volume due to tangential stretch. Moreover, fluid exchange primarily occurs through inter-endfoot gaps, driven by pressure differences, rather than across the aquaporin-4 (AQP4) rich endfoot membrane. PVS stiffness critically modulates these dynamics: increased stiffness of the PVS, for instance, due to vessel pathology or aging, would minimize or even reverse fluid exchange at the gliovascular interface. While AQP4 mediated water movement has a negligible impact on pulsation-driven mechanics, it significantly enhances osmotically driven fluid flow. Overall, our findings elucidate the complex balance of forces governing gliovascular mechanics and suggest that PVS composition strongly influences endfoot-parenchymal fluid exchange.

**Significance:** Perivascular spaces, formed by astrocyte endfeet wrapping the vasculature, are high-conduit pathways for brain fluid flow and clearance. Vascular pulsations drive this flow, but the resulting mechanical interactions at the gliovascular interface remain largely unknown. We introduce a computational model of the solid and fluid mechanics here, using realistic geometries to capture intricate astrocyte morphology at the subcellular level. Our simulations reveal that changes in perivascular composition - associated with aging or neurodegenerative diseases - fundamentally alter mechanical coupling, potentially impeding fluid transport. This work provides a mechanistic framework for understanding brain clearance and constitutes a foundational model for computational studies of mechanical forces in the nervous system.

---

Astrocytes, the main macroglial cell of grey matter, have exceedingly complex morphology. In addition to having tiny cellular protrusions that fill the gaps between the other cells of the neuropil, astrocytes form a sheath around the brain’s vasculature, by endfoot processes extending from the soma towards the vessel, or by residing juxtavascularly (1). Thus, endfeet form a nearly continuous barrier layer around the brain’s vasculature (1, 2) and define the outer perimeter of the perivascular spaces (PVS). However, the precise role of astrocyte endfeet remains enigmatic. In some sharks and other elasmobranches, the endfoot sheath is believed to be the main constituent of the blood-brain barrier (3), whereas in mammals, gaps between neighbouring endfoot processes hinder the endfeet from acting as a tight barrier. Endfoot signaling has been suggested to play a role in blood flow control (4, 5), but these mechanisms remain controversial.

A primary function proposed for this gliovascular architecture is its role in brain waste clearance through the so-called glymphatic system (6). In this system, the PVS allow the cerebrospinal fluid to efficiently distribute through the brain, exit into the tissue, exchange with the interstitial fluid, and thus clear out extracellular waste products, such as beta-amyloid. These mechanisms were shown to be considerably more effective in sleep (7). Due to its role in the removal of extracellular waste products, the glymphatic system is considered a key physiological mechanism preventing neurodegenerative disease. The concept extends foundational work by Helen Cserr and colleagues, demonstrating that PVS enable faster-than-diffusion movement of solutes throughout the brain (8, 9). Although passive vessel pulsations from the heartbeat and respiration as well as active vascular dynamics likely are the main drivers of this extracellular waste clearance system, the mechanical forces and interactions between the blood vessels, the surrounding PVS, and the astrocytic endfeet remain only rudimentarily understood.

Potentially central to this mechanical interplay are aquaporin-4 (AQP4) water channels, which cover nearly 50% of the surface area of the endfeet facing the vasculature. AQP4 is a passive, bi-directional transmembrane water channel, a tetramer that can form supramolecular structures in the form of so-called orthogonal arrays of particles (10). In the brain, AQP4 has been extensively studied in the context of pathophysiology, in particular in brain edema. Here, removal of aquaporin-4 may be protective for certain types of edema formation, but detrimental for other types of edema. Further, AQP4 expression patterns are known to be disrupted in a range of brain disorders, both when studied in animal disease models, or in patient tissue (10). The role of astrocytic AQP4 in physiology, however, remained largely unclear until the proposal of the glymphatic system implicated the channel in brain-wide waste clearance (6). The initial discovery revealed that mice devoid of AQP4 clear interstitial waste more slowly (6), a finding reinforced by follow-up studies demonstrating that the high density of AQP4 at the gliovascular interface is critical for this function (11–14). However, the precise mechanisms by which AQP4 facilitates brain waste clearance stands as an open question.

In addition to the role of membrane permeability, the overall fluid dynamics are also governed by the structural properties of the PVS itself. Recent evidence highlights that the composition of the PVS is heterogeneous between different vessels and may change with aging and vessel pathology (15). Crucially, changes in the PVS’ composition of extracellular matrix proteins, associated with ageing and Alzheimer’s disease and potentially involving dysfunctional perivascular macrophages, have been proposed to affect both vessel dynamics and perivascular fluid dynamics (16). It is beyond existing experimental approaches to probe how altered PVS composition affects mechanical coupling between the vessel, PVS and endfeet.

Understanding this mechanical coupling is critical, as mechanical forces – including fluid flow, mechanical stress, and tissue deformation – are integral to how physical interactions can affect brain function, development, and clearance (17, 18). Despite their importance, quantifying these forces at the cellular level presents significant challenges. While electron microscopy (EM) provides high spatial resolution, as a post-mortem method it cannot capture functional dynamics and introduces significant artifacts. Conversely, in vivo imaging such as two-photon microscopy lacks the resolution required to resolve mechanical forces and fluid movements within the brain’s complex microenvironment. Computational modelling emerges as a powerful tool to overcome these challenges, offering a non-invasive approach to simulate and analyze the biomechanical interactions of neural and glial cells, extracellular matrix and the cerebral vasculature. By harnessing structural imaging techniques, high-performance computing and novel modelling approaches, computational models can provide insights into the impact of mechanical forces on cellular behavior, helping to shed light on complex physiological processes that are otherwise impossible to measure directly.

In this study, we use 3D electron microscopy data to create a poroelastic model of an arteriole segment with surrounding PVS, endfeet and neuropil, and perform high-fidelity computational modeling of the mechanical interplay at the gliovascular interface driven by cardiac-induced arteriolar pulsations. We find that arteriole dilation induces an endfoot sheath expansion, and drives fluid into the surrounding neuropil, but interestingly, almost exclusively through the endfoot gaps rather than through AQP4 in the endfoot membrane. We demonstrate that oscillatory pressure and fluid flow patterns are governed by a complex balance of radial compression and tangential stretch of the structures involved, suggesting that the PVS composition of extracellular matrix and cells that may occur in aging and disease may strongly affect gliovascular dynamics and endfoot–parenchymal fluid exchange. Importantly, we find that PVS stiffening reverses the PVS volume, pressure and flow dynamics across the cardiac cycle, with minimal fluid exchange at certain PVS stiffnesses.

## Results

### Digital high-resolution reconstruction and representation of the astrocyte endfoot sheath

The gliovascular unit, formed by astrocytes and astrocytic endfeet, ensheathing vascular and perivascular spaces, dynamically and rhythmically moves in response to vessel dynamics (Figure 1A-B). In order to create a high-fidelity in silico platform for predicting the mechanical response of this fundamental structure, we turned to a high-resolution segmentation of the extravascular surroundings of an arteriole in the mouse visual cortex (20) (Figure 1C). Along a 20 µm-segment of this vessel, we identified six astrocyte endfeet, together ensheathing a PVS surrounding the vessel lumen. Using the fTetWild meshing software (22), we created a conforming tessellated representation of the individual endfeet, the PVS, and the remaining (non-PVS) ECS (Figure 1D-E), while compensating for tissue shrinkage due to chemical fixation (see Methods). The mean arteriole diameter of 12.2 µm and mean PVS width of 4.6 µm resulted in a mean inner endfoot sheath diameter of 16.8 µm and an outer diameter of 21.0 µm (Figure 1F). In terms of volume, the PVS occupied 41% of the reconstructed geometry, while each endfoot occupied between 0.4 and 9%, giving a total of 30% for all endfoot fragments combined (Figure 1G). With an additional 19% for other neuronal and glial cells next to the endfeet, the ECS occupied the remaining 10%, which translates to an extracellular volume share of 16.6% of the endfoot sheath alone (disregarding the PVS). Characterizing the shape and width of the ECS, we observed a heterogeneous geometry, with tight inter-endfeet gaps and channels (10–50 nm), most spaces ranging between 20 and 120 nm, but also some larger pools up to 500 nm wide (Figure 1H).

**Fig. 1.**
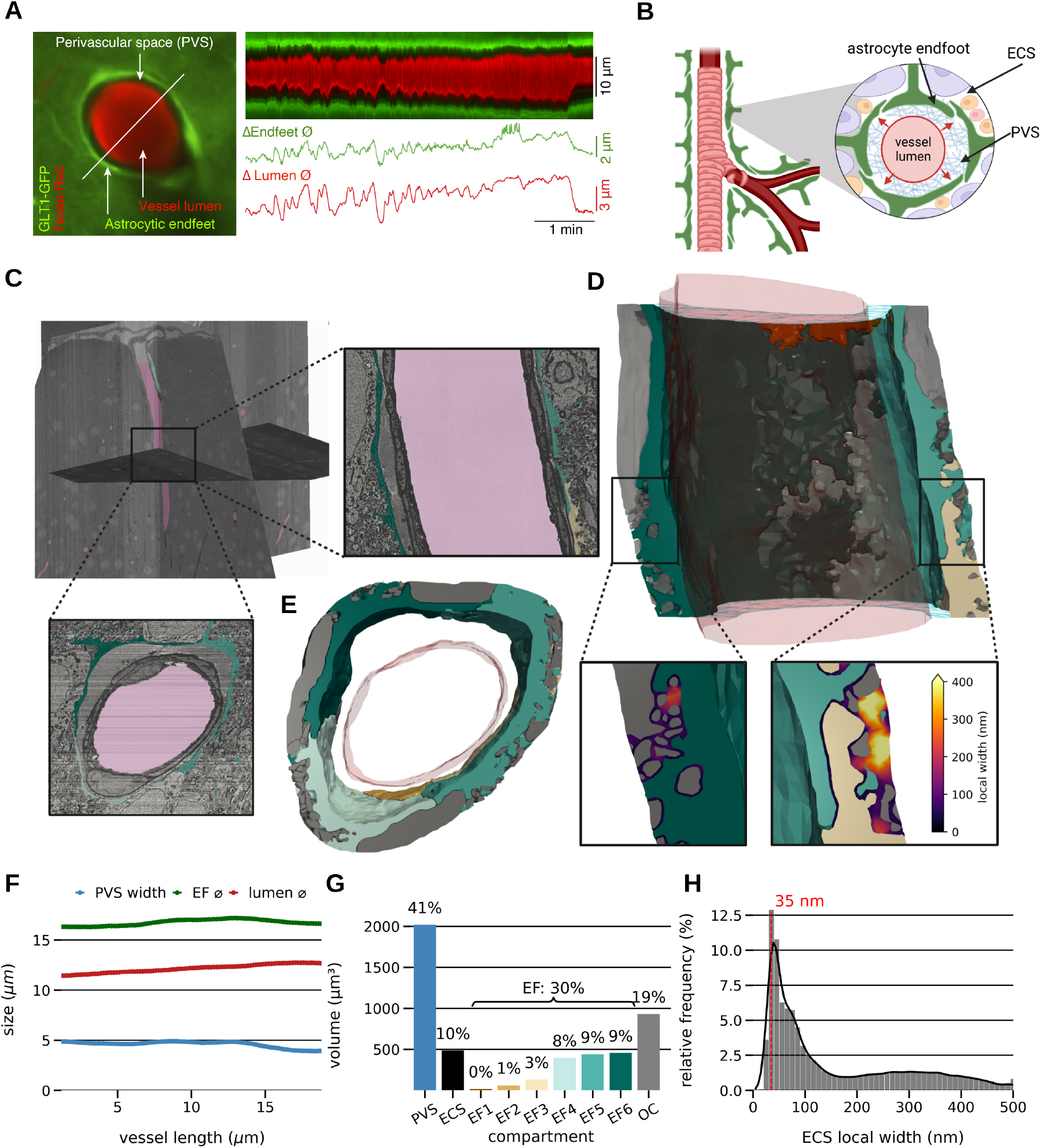
Geometrically-detailed reconstructions of the extravascular cellular environment enable high-fidelity simulations of the gliovascular interplay. A) Two-photon imaging of a penetrating arteriole shows dynamic and substantial changes in endfoot sheath diameter (green) and PVS width (black) following vessel dilation and constriction (vessel lumen in red) (adapted from Bojarskaite et al. 2023 (19)), here and below ∅ denotes diameter; B) sketch of a penetrating arteriole in the mouse visual cortex and its surroundings including the PVS, ECS, and endfeet; C) electron microscopy image overlayed with the astrocyte segmentations (data extracted from the IARPA MICrONS dataset (20), interactive neuroglancer visualization). We remark that the PVS is thought to (at least) partly collapse in postmortem tissue (21). D) The reconstructed geometry (radial cut and zoom-ins shown), with endfeet (brown to blue-green), other cells (grey) and lumen surface (pink). The PVS and ECS are also part of the geometry. The zoom-ins are color-coded by the local width of the ECS to highlight the range of gap widths (see also H); E) Radial cut through the reconstructed geometry (colors as in D); F) average PVS width, inner endfoot sheath diameter and vessel lumen diameter along the longitudinal direction of the arteriole (values corresponding to a radially symmetric geometry of the same cross-sectional area); G) volume of the PVS, endfeet (EF1-6), other cells (OC) and ECS; H) local geometry of the non-PVS ECS, characterized by the distribution of local widths (defined per computational element as the diameter of the largest sphere fitting into the ECS and containing the element midpoint).

We then represented the spatially-resolved astrocyte endfeet and other neural and glial cells, the PVS and the ECS as separate domains and considered each as deformable and permeable (poroelastic) structures, separated by the semi-permeable cell membranes. We set the stiffness of the astrocyte endfoot skeleton to be 10× that of the ECS and PVS, and the permeability of the PVS to be 10× that of the ECS, which in turn is set twice that of the astrocyte endfeet. Crucially, we also distinguished between the permeability of the AQP4-rich adluminal endfoot membrane and the abluminal side (see Methods). The mechanical response of this environment is then described by the displacement field **d** and the fluid pressure field *p*, with the fluid flux **q** and stress tensor ***σ*** as auxiliary derived fields, varying across space and time.

### Stretch dominates squeeze: arteriole dilation compresses the PVS while expanding the endfoot sheath

This gliovascular environment responds to external or internal forces by distributing and balancing elastic forces and fluid motion, in a manner that depends on a number of factors (morphology, stiffness, permeability) and to an extent that is hard to predict a priori. Therefore, to investigate the dynamics associated with typical heartbeat-derived vascular pulsations, we prescribed a 10 Hz-pulsatile dilation and contraction of the inner arteriole surface (± 0.8%, (23)) and analyzed the resulting computational predictions of movement and force distribution within the PVS and endfoot sheath.

As expected, the dilation led the entire structure to expand (Video SV1), nearly uniformly along the vessel length and synchronously, but with decreasing displacement magnitude towards the outer rim of the endfoot sheath. The mean diameter change ranged from 0.1 µm (vessel lumen) to 0.06 µm (inner endfoot sheath) at peak systole (Figure 2A). Notably, the PVS itself decreased, both in width (− 0.01 µm) and volume (− 0.03%) (Figure 2A–C). In contrast, the astrocyte endfoot volume increased slightly (+0.001%), while the ECS volume increased (0.13%) (Figure 2C). The largest local increases in ECS volume occurred in the vicinity of the tight inter-endfeet gaps (up to 0.6%, Figure 2D). Thus, in terms of overall volumes, the compression of the endfoot sheath due to arteriole dilation was overcompensated by the stretch induced by its increased radius.

**Fig. 2.**
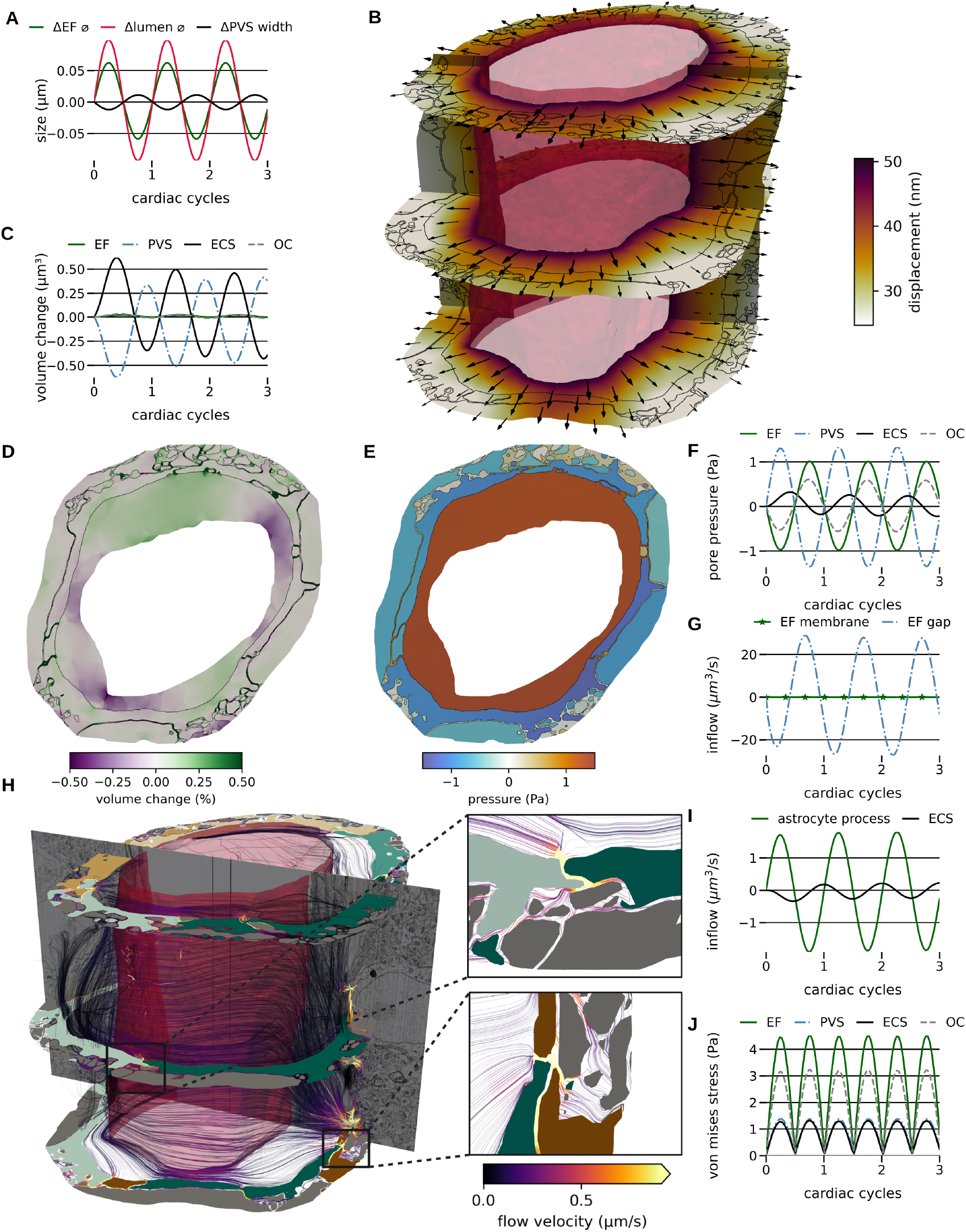
Stretch dominates squeeze: endfoot sheath response to cardiovascular pulsations. A) Change of mean lumen diameter, endfoot sheath diameter, and PVS width over three cardiac cycles, ∅ denotes diameter; B) displacement magnitude at peak arterial dilation: cross-sectional and longitudinal slices color-coded by displacement magnitude, arrows indicating displacement direction, cell membranes outlined in black, vessel lumen in transparent red; C) volume change of endfeet (EF), PVS, ECS and other cells (OC) over time; D) volume change at peak arterial dilation (cross-section); E) fluid pressures at peak arterial dilation (cross-section); F) mean fluid pressures of the endfeet, PVS, ECS and other cells over time; G) flow rates *into* the PVS via the adluminal endfoot membrane (solid with stars, green) and through the endfoot gaps i. e. the interface between PVS and endfoot sheath ECS) (dot-dashed, blue) over time; H) flow streamlines in the PVS and ECS at peak systole with cross-sections through the endfeet and other cells (colors as in the Figure 1) and vessel lumen (transparent red); flow streamlines colored by the velocity magnitude; zoom-ins illustrate rapid flow through the endfeet gaps. I) Flow in through the outer radial boundaries: inflow rates for the astrocyte processes (AP, outer boundary of the astrocyte endfeet) and the ECS outer boundary; J) mean von Mises stress of endfeet, PVS, ECS and other cells.

The mean fluid pressure in the PVS and in the ECS increased by up to 1.3 Pa and 0.3 Pa, respectively, with the PVS fluid pressure peaking in sync with the peak arteriole dilation (Figure 2E–F, Video SV2). On the other hand, the pressure within the astrocytic endfeet dropped by -1.0 Pa. This seemingly counter-intuitive pressure reduction at peak arteriole dilation is a direct consequence of the small increase in endfoot volume. In turn, these pressure differences induced fluid exchange back and forth between the PVS and the endfoot sheath over time. Interestingly, the flow rates through the endfoot gaps strongly dominated those through the adluminal endfoot membrane, peaking at 29 µm3/s versus at 0.01 µm3/s, with peak flow velocities in the endfeet gaps of about 0.8 µm/s (Figure 2G–H). In addition, the fluid exchanged between the endfoot sheath and the outer surroundings. The drop in endfoot fluid pressure during arteriole dilation pulled fluid in at rates up to 1.8 µm3/s through the continuation of the astrocyte process. Simultaneously, we observed flow, though of lower magnitude and opposite directionality, through the ECS: fluid moves to and from the surrounding tissue via the outer extracellular surface at rates up to 0.23 µm3/s (Figure 2I). Thus, most of the fluid crossing the endfeet gap contributes to an increase of the ECS volume within the endfoot sheath and does not penetrate further into the surrounding tissue. Examining the mechanical stresses, we found the highest mean von Mises stress in the astrocyte endfeet (up to 4.5 Pa at peak systole) and substantially lower stress on the PVS and ECS (1.4 Pa and 1.3 Pa peak mean stress, respectively, Figure 2J).

### PVS stiffening reverses gliovascular flow dynamics

PVS morphology and composition change with aging and disease, potentially due to perturbed parenchymal macrophage activity or plaque deposition (15, 16). Assuming that such structural changes modify the elastic properties of the PVS, we next asked how stiffness affects the mechanical interaction between the vascular pulsations, the PVS and endfoot sheath (Figure 3A).

**Fig. 3.**
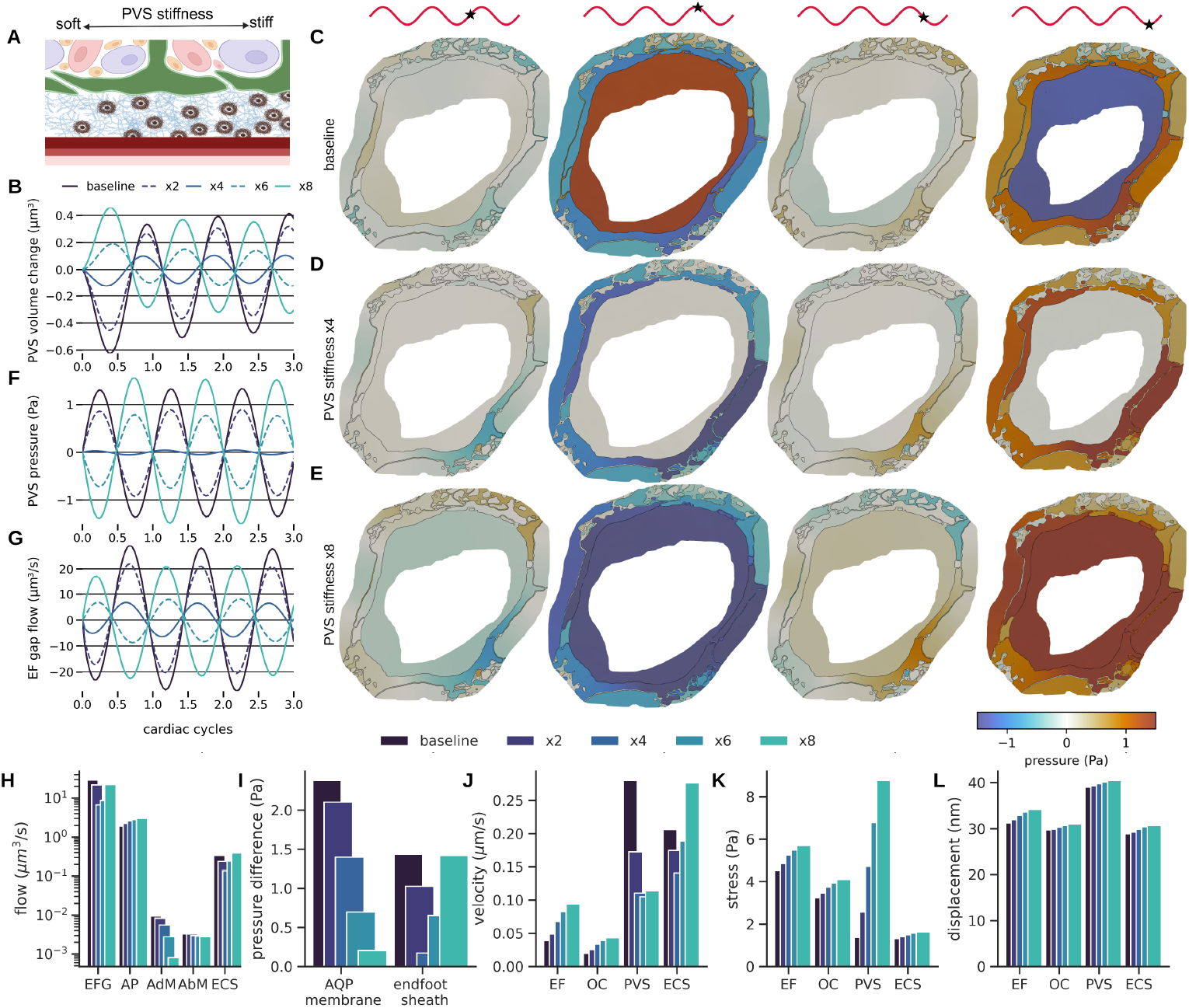
PVS stiffening reverses gliovascular flow dynamics. A) Conceptual illustration of PVS stiffening due to e.g. plaque accumulation or changes in the extracellular matrix composition; B) change in mean PVS volume over time for models with an increase in PVS stiffness by a factor of 1, 2, 4, 6, and 8. C–E) cross-section of the geometry, showing the fluid pressure over one cardiac cycle for the baseline model (C), PVS stiffness increased by 4× (D) and 8× (E); F) mean PVS pressure over time; G) flow through the endfoot gaps over time; H) peak fluid flow rates across the endfoot gaps (EFG, interface between PVS and ECS), the outer astrocyte process (AP, outer radial model boundary of the endfeet), the adluminal endfoot membrane (AdM), the abluminal endfoot membrane (AbM) and the ECS outer radial boundary (ECS); note the logarithmic scale. I) Peak mean pressure gradient across the adluminal endfoot membrane and the whole endfoot sheath; J) peak mean flow velocities in each compartment; K) peak mean von Mises stress in each compartment; L) peak mean displacement magnitude in each compartment.

Interestingly, PVS stiffening transforms PVS volume dynamics in response to cardiac-driven vascular pulsations. While we had until now observed a decrease in PVS volume in response to the vessel dilation, the behavior was reversed by an increase in PVS stiffness: an 8 × stiffer PVS yields a volume *increase* of similar magnitude as the previously observed volume *decrease* (Figure 3B). We attribute this phenomenon to a change in the relative strength of two effects. On the one hand, the PVS is radially compressed due to vessel dilation; on the other hand, the outer PVS is tangentially stretched due to an increase in PVS radius. Stiffening reduces the compression, and thus outer stretch dominates at vessel dilation. Moreover, this also implies that at a certain PVS stiffness, stretch and compression balance each other, leading to an almost constant PVS volume. Our simulation results indicate that this occurs at approximately a 4 × increase in PVS stiffness compared to our baseline (Figure 3B).

These changes in volume dynamics in turn induced a reversal in pressure and flow behavior. Comparing with the baseline PVS pressure increase of 1.3 Pa at peak dilation, we found a comparable pressure decrease with an 8 × stiffer PVS, and almost constant PVS pressure over time at 4 × increased PVS stiffness (Figure 3C–F). We also observed substantial changes in the endfoot gap flow. In our baseline model, fluid flows from the PVS through the endfoot gaps into the ECS at peak vessel dilation, while higher PVS stiffnesses yield flow into the PVS (Figure 3G–H). In the scenario with a 4 × increase in PVS stiffness, the fluid exchange between the PVS and the parenchyma was substantially reduced. These effects on gliovascular fluid exchange would likely have a profound impact on solute exchange between the PVS and surrounding tissue driven by oscillatory flow and mixing.

In addition to the effect of PVS stiffness on fluid flow rates (Figure 3G–H), we systematically investigated the effect of PVS stiffening on the pressure differences, mean fluid velocities, stress and displacement magnitudes (Figure 3I–L). Our analysis showed that the pressure difference across the adluminal endfoot membrane decreased with increasing PVS stiffness, and consequently also the resulting peak flow rates. In contrast, the peak pressure difference across the complete endfoot sheath depends nonlinearly on the PVS stiffness; dropping from 1 × to 2 × to 4 × but then increasing from 4 × to × 6 to 8 ×, and in fact reaching the same magnitude at 8× stiffer PVS (but with the opposite phase) (Figure 3I). In terms of mean fluid velocities, PVS stiffening led to a moderate increase in intracellular flow (in the endfeet and other cells), while reducing PVS flow velocities by more than a factor 2 (Figure 3J). As for the endfoot gap flow rates and endfoot sheath pressure gradients, the mean flow velocities in the ECS first dropped, but then increased and even exceeded the baseline model as the PVS stiffness increased (Figure 3J). Finally, PVS stiffening increased stress on the solid components of all compartments, most prominently in the PVS, while the peak mean displacement rose only slightly (Figure 3K–L).

### Robustness and sensitivity to perivascular pathway resistance and waveforms

Considering that the resistance of the perivascular pathway is subject to substantial debate (24–31), a natural question is to what extent the previous observations depend on the flow features within the PVS. There, we considered the extreme case of a *maximal* perivascular pathway resistance, corresponding to fluid moving primarily radially within and from the PVS, by prescribing zero flow at both ends (front/rear) of the PVS. To also consider the other extreme, a perivascular pathway with *minimal* resistance, we allow the fluid to move freely in and out of the PVS at the rear end, with parameters otherwise as at baseline (Figure S1A).

The two scenarios induce clearly different PVS flow patterns: the minimal resistance scenario resulted in oscillatory axial flow with linearly increasing velocities towards the rear end reaching up to 2 µm/s, while we found a slower, more heterogeneous flow pattern converging towards the endfeet gaps in the maximal resistance scenario (Figure S1B–C). The lower resistance also resulted in lower PVS pressures (peaking at 1.0 Pa vs 1.3 Pa). Conversely, radial CSF flow rates and fluxes were reduced in the low resistance scenario, most strikingly through the endfoot gaps (-43%) (Figure S1D–E). The reduced radial flow was reflected by smaller pressure differences across the endfoot membrane (-32%) and across the whole endfoot sheath (-41%) (Figure S1F). While the mean fluid velocity in the PVS rose by more than five-fold, the flow in the ECS and within the endfeet slowed down by -30% and -10%, respectively (Figure S1G). The reduced resistance was also accompanied by smaller stresses in the endfeet and extracellular matrix, larger stress in the PVS, and smaller displacements overall (on the order of ± 10%) (Figure S1H–I). Examining the overall diameter changes of the endfoot sheath, the PVS width and the synchronicity of the system, we found a minor reduction in endfoot diameter changes and an increase of PVS width changes with lower resistance, and endfoot and vessel movement remained in synchrony.

In addition, we assessed whether the waveform of the arterial pulsation influenced the results. Physiological pulsations are often asymmetric, characterized by a steep dilation phase followed by a slower constriction. We performed simulations using a physiological, asymmetric waveform extracted from experimental data (23). To distinguish the effects of asymmetry from amplitude, we scaled the waveform to match either the maximum amplitude or the area under the curve (AUC) of our baseline sinusoidal model (Figure S2). We found that waveform asymmetry had a negligible effect on the overall gliovascular mechanics (Figure S3). While the steeper expansion phase caused a transient increase in peak flow velocities (up to a factor of two in the AUC-scaled case), the integrated flow volumes, stress distributions, and pressure gradients remained comparable to the symmetric baseline.

Finally, we remark that, in contrast to the maximal resistance model, the low resistance simulation results depend on the length of the vessel segment, and with increasing segment length, the cumulative resistance increases. Thus, the maximal resistance model can be interpreted as a long-segment limit of the minimal resistance model. We conclude that the maximal resistance model is the more relevant case here, and consider the deviations in radial motion (between 7 and 58%) as upper bounds of the error of this assumption.

### Low-frequency, high-amplitude vasomotion drives large tissue displacement and fluid exchange

To investigate the effects of neurovascular coupling on gliovascular mechanics, we also simulated a “vasomotion” scenario characterized by low-frequency (0.2 Hz), high-amplitude (8% dilation) vessel wall motion, mimicking the functional hyperemia observed in response to whisker stimulation (32) or sleep-state dependent vasomotion (19). Consistent with the large (10-fold) increase in arterial dilation amplitude, our model predicts a substantial increase in the mechanical response of the surrounding tissue compared to the baseline “cardiac” scenario (10 Hz, 0.8% dilation). Peak displacement magnitudes increased by 877% in the endfeet and 900% in the ECS compared to the cardiac baseline (Figure S5A). Counter-intuitively, this large increase in displacement did not uniformly amplify fluid velocities. The flow rate through the inter-endfoot gaps (EFG) actually decreased by 8% (Figure S5A). The disparity is explained by the frequency-dependent poroelastic response of the tissue: the slower dilation of the vasomotion regime (0.2 Hz) allows sufficient time for pressure equilibration. Consequently, the PVS and ECS pressures become almost synchronized, preventing the formation of large gradients across the endfoot layer and limiting the amplification of local flow velocities (Figure S5B–C). However, the absolute pressure amplitude rises substantially during vasomotion. This global pressurization drives increased exchange across high-resistance pathways. Transmembrane fluxes (across AdM and AbM) increased by 696– 1453%, and most prominently, the fluid efflux into the surrounding tissue increased by 3288%, due to the combined effect of larger absolute pressure changes, and pressure synchronization between PVS and ECS (Figure S5A).

### Osmotic challenge drives AQP4-enhanced net fluid flow

Blood osmolytes such as glucose have been suggested as potential drivers of brain fluid flow (33), with reported persistent concentration differences between vascular and extracellular glucose amounting to 4 mM (34). Noting that the constant supply of glucose requires a spatial concentration gradient across the gliovascular interface, we hypothesized that perivascular glucose might be elevated compared to the surrounding tissue, and more specifically, glucose levels might be elevated compared to the astrocyte endfeet. Here, we investigated whether and to what extent adding osmolytes in the gaps between the endfoot and vessel could drive fluid flow in the model, and how the presence of AQP4 water channels affects gliovascular mechanics under such conditions. Concretely, we considered a scenario without vascular pulsations but prescribed an osmotic pressure gradient of 1 kPa - corresponding to an osmolyte concentration difference of 0.39 mM - across the endfoot membrane facing the blood vessel as an alternative driving force (Figure 4A).

**Fig. 4.**
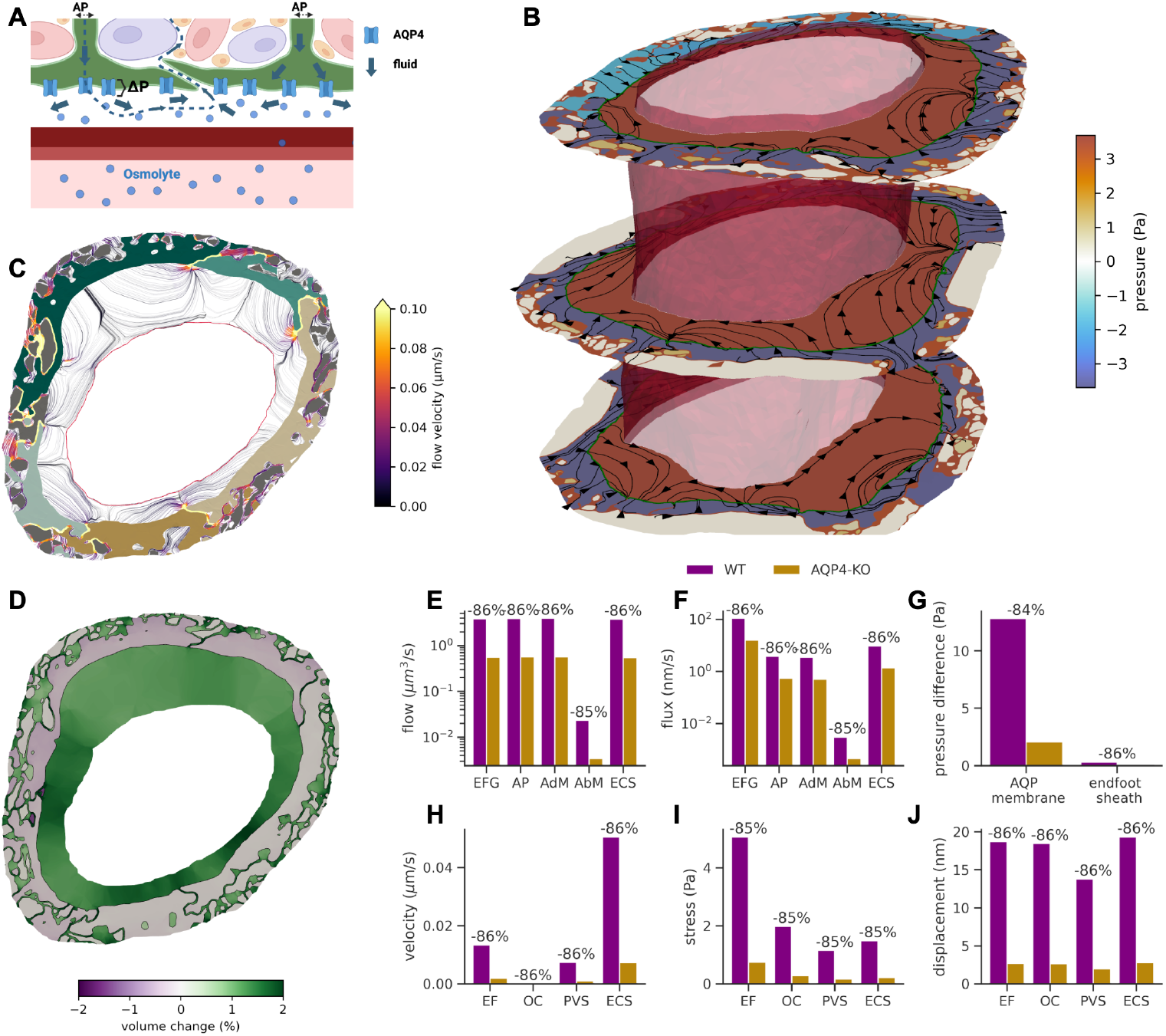
Osmotic differences can drive fluid exchange between the PVS and its surrounding tissue that depends on AQP4 density. A) A blood osmolyte such as glucose permeates the PVS, potentially inducing an osmotic pressure difference across the endfoot membrane facing the vessel, yielding a persistent flow through the endfoot, into the PVS and further through the endfoot gaps into the surrounding tissue. B) The fluid pressure and flow streamlines in the endfeet, ECS and PVS (cross-section) induced by the osmotic pressure difference; C) extracellular fluid flow velocities (streamlines colored by velocity magnitude); D) relative volume change; E) mean fluid flow rates across the endfoot gaps (EFG, interface between PVS and ECS), the outer astrocyte process (AP, outer radial boundary of the endfeet), the adluminal endfoot membrane (AdM), the abluminal endfoot membrane (AbM) and the ECS outer radial boundary (ECS); F) mean flux rates (labels as in E); note the logarithmic scale in E) and F). G) Mean pressure gradient across the adluminal endfoot (AQP) membrane and the whole endfoot sheath; H) flow velocities in each compartment; I) mean von Mises stress in each compartment; J) mean displacement magnitude in each compartment.

The osmotic pressure gradient induced a steady, swirling flow pattern, with a flow rate of 3.8 µm3/s across the endfoot membrane leading to a pressure increase of about 4 Pa in the PVS (Figure 4B). The intracellular fluid pressure dropped by up to 5 Pa, with considerable variations between the endfeet depending on their surface area exposed to the osmotic gradient. The resulting pressure gradients induced both intra- and extracellular net fluid flow: the intracellular pressure drop pulled fluid from the surrounding tissue via the intracellular pathway into the endfoot, while the increased PVS fluid pressure drove interstitial fluid through the endfoot gaps into the extracellular space and further into the surrounding tissue, creating a persistent mechanism for fluid exchange between the PVS and the surrounding cerebral tissue (see conceptual illustration in Figure 4A). Peak endfoot gap flow velocities reached up to 100 nm/s in this scenario (Figure 4C). Additionally, the elevated PVS pressure also induced an increase in PVS volume by up to 2%, while endfoot volumes decreased slightly (Figure 4D). We also simulated this scenario with a reduced PVS permeability (comparable to the ECS), giving similar results.

Finally, we asked how the water permeability of the astrocytic endfoot membrane would affect this osmotically-induced fluid flow. Again inspired by the reduced water permeability measured in AQP4 knock-out mice (35), we decreased the adluminal membrane permeability by 7 × before repeating the simulation. In this scenario, the seven-fold reduction in membrane permeability directly translated into a seven-fold reduction in transmembrane fluid flow (−86%), and subsequently the same relative reduction in flow through the astrocyte processes, endfeet gaps, and ECS (Figure 4E–F). Indeed, we observed an equal relative reduction in all other quantities of interest, including the pressure difference across the adluminal membrane and endfoot sheath, mean fluid velocities, stress and mean displacement in each compartment (Figure 4G–J). Although this modelling scenario is based on assumptions that are not possible to experimentally verify with current technology, it illustrates how osmotic driving forces at the gliovascular interface may be a potent driver of directional fluid flow on the microscale, and that AQP4 may play a key role in this mechanism.

## Discussion

We have presented a detailed computational model of solid and fluid mechanics in astrocyte endfeet at the gliovascular interface using realistic high-resolution geometries derived from electron microscopy. A key model prediction is that during arteriole dilation, the PVS is compressed while the endfoot sheath expands. Under both cardiac and vasomotion pulsations, this leads to increased fluid pressure in the PVS and, somewhat surprisingly, decreased pressure within the endfeet. During the high-amplitude wall motion of vasomotion, tissue deformations are substantially larger, driving substantial fluid volume exchange with the surrounding parenchyma by sustaining flow over longer periods. Another important prediction for understanding extracellular fluid dynamics in aging and pathology is that increasing PVS stiffness reverses the PVS volume, pressure and flow dynamics across the cardiac cycle. In some cases, this results in minimal fluid exchange between the PVS and surrounding tissue. Hence, the cellular and extracellular matrix composition of the PVS may strongly affect gliovascular dynamics and endfoot–parenchymal fluid exchange. We also found that the resistance of the downstream perivascular pathway influences local flow patterns: lower resistance leads to reduced radial flow and increased axial flow within the PVS, but only minor changes in endfoot stress and displacement. Moreover, fluid leaves the PVS predominantly through the inter-endfeet gaps rather than through the AQP4-rich endfoot membrane. Consequently, even substantially reducing the AQP4-mediated water permeability had a negligible impact on the dynamics driven by vascular pulsations – despite a considerable relative decrease in fluid flow across the endfoot membrane. In contrast, when driven by an osmotic pressure gradient, an AQP4-rich adluminal endfoot membrane enhances fluid exchange between the PVS and surrounding tissue.

While both oscillatory and directional net flow of CSF along the PVS have been extensively studied in both animal (16, 23, 32) and mathematical models (24–27, 29– 31), very few studies realistically address fluid exchange at the gliovascular interface and the dynamic mechanical interplay between the blood vessel, the PVS and the endfeet. Recent data clearly demonstrate that astrocyte endfeet move with blood vessel dynamics, especially during sleep (19), but the outer boundary of the PVS and the endfeet themselves are typically modelled as a rigid sheath. A notable exception is the study by Kedarasetti et al. (36), in which the authors also modelled a poroelastic (but homogenized) tissue surrounding the PVS. Koch et al.(37) relied on a non-deformable and idealized geometry, but explicitly accounted for endfoot gaps — with sizes and distributions aligning with literature — to theoretically quantify endfoot sheath permeability. To the best of our knowledge, ours is the first computational model of the gliovascular interface using detailed, realistic geometries that capture the complex morphology of the astrocyte endfoot sheath, and thus enable high-fidelity simulations of flow through the endfoot membrane and the inter-endfoot gaps. Our poroelastic modelling approach describes the complex interaction of mechanical forces, deformation and fluid flow – both within extra- and intracellular spaces and across cell membranes – and thus constitutes a foundational model for computational studies of mechanical forces in the nervous system. In the light of the emerging role of mechanobiology in neuroscience (17, 18, 38), our model establishes a starting point for future in silico studies of cellular mechanics, especially in combination with dense neuronal tissue reconstructions.

Turning to model validation, we first note that due to the complexity of measuring many of the relevant quantities in vivo, several of our computational predictions lack experimental counterparts. However, both Holstein et al. (32) and Bojarskaite et al. (19) report that arterial dilation by whisker-stimulation-induced hyperemia and sleep state changes directly affect the PVS width in head-fixed mice. They describe that arterial dilation induces a dilation of the endfoot sheath, but with a lower magnitude, thus reducing the width of the ensheathed PVS. This aligns well with our findings. Regarding tissue deformation, our vasomotion simulations predict substantial motion, resulting in peak tissue displacements of approximately 300–400 nm. This aligns in order of magnitude with the 0.8–1.0 *µ*m radial displacement of astrocytes reported experimentally during whisker-stimulation-induced hyperemia (32). The higher values observed in vivo likely reflect the larger vessel diameters (≈ 58 *µ*m MCA) and wall displacements in the experimental measurements compared to our modeled penetrating arteriole. Moreover, Holstein et al. (32) report an increase of both radial and axial PVS flow velocity after whisker stimulation, with radial velocities in the order of 0–4 µm/s. Our predicted peak mean radial PVS flow velocities (0.07– 0.28 µm/s) and local maxima (approximately 1 µm/s) fall well within this experimental range. In terms of intracellular cytosolic flow rates, our estimates (up to 0.03 µm/s peak mean velocity in the endfeet) align well with typical values for cytosolic flow rates of approximately 0.01–1 µm/s (39) and recent observations of deformation-induced cytosolic flows (40).

We are not aware of any reports on the mechanical stress in astrocyte endfeet during physiological pulsations. However, our computational predictions of up to 5 Pa (cardiac) and 42 Pa (vasomotion) fall at the lower end of typical stresses in the cytoskeletal mesh, which can sustain hundreds of pascals (39, 40). This suggests that physiological wall motion—even in the high-amplitude vasomotion regime—generates low to moderate mechanical loads compared to the structural limits of the astrocyte cytoskeleton. Furthermore, assessing our transmembrane flux rates ranging around 10−2 nm/s in the cardiac-induced scenario and around 10 nm/s in the osmotically-driven scenario, we find good agreement with characteristic flow rates across membranes of 1–10 nm/s at characteristic pressures of 100 Pa (39), if the difference in transmembrane pressure gradients is taken into account.

The composition and morphology of the PVS changes with aging and in disease, potentially due to accumulation of extracellular matrix proteins or plaque deposition (15, 16). Investigating how changes in PVS elastic properties affect gliovascular mechanics, we found that PVS stiffening reverses the flow dynamics between the PVS and its surrounding tissue: a soft PVS compresses with vessel dilation and pushes fluid through the endfoot gaps, while a substantially stiffer PVS experiences a larger tangential stretch than radial compression and draws fluid through the endfoot gap. While these two cases lead to oscillatory fluid exchange with similar magnitude and opposing phase between PVS and surrounding tissue, an interesting third case emerges: at a certain PVS stiffness, radial compression and tangential stretch of the PVS balance each other, resulting in (almost) no fluid exchange across the endfoot sheath. This scenario arose at about a four-fold increase in PVS stiffness compared to our baseline model, providing a mechanistic explanation for reduced glymphatic flow with PVS plaque accumulation and perivascular stiffening.

In this study, we used a linear poroelasticity model, in part to avoid additional assumptions relating to (uncertain) nonlinear material characteristics. Our baseline cardiac pulsation scenario induces vessel wall displacements of approximately 0.8% and tissue strains well within the generally accepted limit for infinitesimal strain theory (***ε*** ⩽ 1%); we thus consider the linear theory to be well-justified here. A fundamental property of this linear framework is that zero-mean oscillatory motion results in a symmetric hydraulic response where the time-averaged flux vanishes (41). Consistent with this principle, a temporally asymmetric waveform did not yield directional bulk flow in our simulations. This contrasts with the study by Kedarasetti et al. (36), which reported net flow driven by asymmetric waveforms; however, their model incorporated non-linearities via nonlinear constitutive laws and large deformations, effectively breaking the symmetry between the dilation and constriction phases. Thus, temporal waveform asymmetry alone is insufficient to drive bulk flow in a linear regime.

Instead, significant net transport requires either substantial local non-linearities such as deformation-dependent material properties (41) or valve-like geometric mechanisms (42) — or driving forces that emerge on a larger spatial scale. Indeed, whether endfeet could act as one-way valves for interstitial fluid flow by changing their gap size in response to pressure changes, as recently proposed (42, 43), stands as an interesting hypothesis. Our results show that the ECS volume between the endfeet indeed increases in response to cardiac-induced arterial dilation, albeit only by approximately 0.1%. Whether this would translate to a sufficient reduction in flow resistance and enable a valve-like effect of the endfoot sheath remains unclear. If such local non-linearities are absent, PVS flow must be driven by macro-scale interactions, such as brain movements associated with vascular dynamics and respiration (44–46), non-reciprocal movements of longer vessel segments (30), or the turnover of CSF and metabolic water (47). While such larger-scale effects are beyond the scope of this study, we cannot completely ignore the role of axial PVS flow, as it also impacts radial mechanics. Given the substantial uncertainty about PVS properties in general and resistance in particular, we compared extreme cases and found that the minimal downstream PVS resistance reduced radial flow and tissue forces by only 7–58% compared with a maximal resistance scenario. Interpreting the maximal resistance scenario as the model’s long-segment limit, we consider these ranges as upper error limits of the simplifying assumption of our choice of the maximal resistance model.

The physiological role of AQP4 water channels has been the subject of considerable research activities over the last decade (2, 10), and AQP4 has been suggested to support glymphatic clearance by facilitating water exchange across the endfoot sheath (21). However, our results indicate that AQP4-mediated water permeability has a negligible impact on the mechanical interaction between the vessel and endfeet. This is primarily because the hydraulic resistance of the inter-endfoot gaps is orders of magnitude lower than that of the membrane, regardless of AQP4 expression. This was also the case when considering larger vascular dynamics similar to what is observed during sleep vasomotion and neurovascular coupling. These findings align with previous estimates of the endfoot sheath permeability (37, 48, 49) but do not provide a satisfactory answer to the question of the physiological role of AQP4, especially given the notable abundance of AQP4 channels, and the impact of AQP4 knockout in animal models (21).

Therefore, drawing on earlier experimental findings demonstrating the importance of AQP4 in influencing osmotically driven water influx, specifically in brain edema (50–52), we asked under which physiological conditions AQP4-mediated water flux could play a more decisive role. Concretely, we hypothesized that perivascular glucose might be elevated compared to the adjacent tissue (and specifically the endfoot) and imposed an osmotic pressure gradient across the adluminal endfoot membrane. In such a scenario, the difference in osmolarity creates a circular flow pattern: intracellular pressure differences drive fluid from the astrocyte cell body to the endfoot membrane, where it crosses into the PVS, leading to an increase in PVS volume and pressure, which consequently drives fluid through the endfoot gaps into the surrounding ECS. Interestingly, we observed that this mechanism is highly sensitive to the endfoot membrane permeability: a sevenfold reduction in permeability – mimicking the absence of AQP4 channels – reduced flow sevenfold. These findings suggest that AQP4 water channels’ primary role is in facilitating osmotically, not hydrostatically driven fluid exchange. While the idea that blood osmolytes might drive fluid flow across the gliovascular interface has been suggested previously (33, 48), the existence of such a gradient is far from certain. However, well-established differences between blood and brain ECS glucose levels (34), and the necessity of a constant supply of glucose across the blood-brain-barrier, suggest a glucose concentration gradient across the gliovascular interface and potentially also at the adluminal endfoot membrane. In addition, we remark that our modelling is independent of the specific cause of osmotic imbalance, and remains relevant for other scenarios involving osmotic pressure differences at the gliovascular interface. Due to the linearity of the model, flow rates scale with the imposed osmotic gradient. We note that our model imposes a static osmotic pressure gradient, representing a persistent concentration difference maintained by continuous vascular supply (e.g., glucose influx). We did not simulate the concurrent advection-diffusion of osmolytes into the ECS. In a physiological setting, the transport of osmolytes would likely reduce the effective transmembrane gradient and potentially generate opposing osmotic forces across the astrocytic-ECS interface. Consequently, our predicted flow rates represent an upper bound of osmotically driven flow resulting from a given initial concentration difference.

A natural question is how our findings contribute to current understanding of the glymphatic system and related pathways. The glymphatic theory today highlights three components: periarterial CSF influx, interstitial solute movement, and solute efflux along perivenous spaces, cranial and spinal nerves (28). To summarize the discussion above, our study focuses on one component of periarterial CSF flux, namely the fluid exchange between the periarteriolar space and its outer surroundings including the role of the astrocytic water channel AQP4 in this exchange. We find that the water permeability of the PVS-facing endfoot membrane, which is modulated by the presence of AQP4 channels, has little if any effect on the microscale gliovascular dynamics of flow driven by cardiac pulsations or vasomotion. Instead, we suggest that AQP4 will play a significant role in fluid exchange between the PVS, astrocyte endfeet and ECS driven by an osmotic pressure difference across the astrocytic endfoot membrane. While we do not simulate transport of osmolytes or other solutes here, substantial differences in net flow of CSF/ISF – such as these associated with the presence or lack of AQP4 – would clearly also affect solute transport and clearance.

The high spatial resolution of electron microscopy allowed us to create a highly detailed reconstruction of the astrocyte endfeet, but also comes with limitations, primarily the shrinkage of ECS due to chemical fixation. In particular, the inter-endfeet gap width is crucial for the endfoot sheath permeability, yet has not been accurately measured in living tissue. After processing the cell segmentation with our processing pipeline (see Methods), the distribution of ECS local width peaked at about 35 nm corresponding to the endfoot gap size, which is only slightly larger than the value of 20 nm reported by Mathiisen et al. (2) (also based on EM imaging). While a potential overestimation of endfoot gap size favors flow through endfoot gaps over transmembrane flow, we note that it does not fundamentally change the flow ratio of more than three orders of magnitude. Similarly, the local width of ECS beyond the endfeet gaps in our geometry is heavily influenced by our image-processing pipeline. However, the advent of new imaging technologies such as super-resolution shadow imaging (SUSHI) (53) has enabled accurate characterization of the local geometry of the ECS. Here, we found that the local ECS width in our reconstructed geometry aligns well with the observations of a highly heterogeneous ECS, ranging from small gaps between 10–20 nm to larger pools of 200–500 nm (54). In terms of the PVS geometry, we chose to identify the entire space between the vessel lumen and the astrocyte endfeet as PVS, which reflects the viewpoint of two-photon imaging and resulted in a PVS width of approximately 5 µm, which is in good agreement with previously reported values (19).

In terms of limitations beyond the uncertainty about geometrical features of the PVS and endfoot sheath, one of our primary modelling constraints is the assumption of linear poroelastic media for the intra- and extracellular space. While being a good first-order approximation, it is well known that biological tissue is often characterized by nonlinear stress-strain relations with memory and elements of plasticity (39). For instance, the collagen networks that are a major component of the extracellular matrix can be substantially stiffer under tension than compression, and rather buckle than compress under compressive loading (55). However, accounting for the inherent nonlinear mechanics requires non-existent detailed knowledge of the material behavior to parameterize and advantageously employ such a model. Furthermore, while our baseline analysis utilizes an idealized sinusoidal waveform, we verified these predictions against a physiological, asymmetric waveform. We observed that the waveform asymmetry had a minimal impact on the resulting gliovascular dynamics, supporting our use of the idealized representation.

In conclusion, our model is the first to combine a comprehensive mathematical model of the poroelastic behavior of intra- and extracellular space with a realistic, 3D electron micrograph-derived geometry of the gliovascular interface. The model provides detailed insights into the mechanics of blood vessels, astrocyte endfeet and their ensheathed PVS, and enables accurate predictions of their interplay. These findings supplement and extend insights from experimental studies on the role of astrocyte endfeet in health and disease, and provide a foundation for further computational studies of cellular mechanical interaction in the nervous system.

## Materials and Methods

### Image processing and computational geometry pipeline

Using imaging data from the EARP MICrONS dataset (20), we extracted a segmentation of the extravascular surroundings of a cortical arteriole, including several astrocyte endfeet, a perivascular space surrounding the lumen and endothelial cells, and other ECSs (Figure 1C). We then generated a computational mesh of the geometry, conforming to the interior interfaces, via a custom image processing and meshing pipeline based on the fTetWild meshing software (22). In particular, as part of the preprocessing, we inflated the computational geometry representation of the ECS of the endfoot sheath to 16.6% to counteract the effect of chemical fixation. The computational mesh includes 258 776 vertices and 1 425 200 cells with a mesh resolution of 0.36 ± 0.22 µm.

### Poroelasticity model including material parameters

We characterize both the intracellular (astrocyte endfeet, neurons and other cells) and extracellular spaces as poroelastic structures composed of an elastic solid matrix filled with cytosol and interstitial fluid (ISF), respectively (39, 56, 57), though with differing material parameters. In the intracellular space Ω_*i*_, this elastic matrix represents the cytoskeleton, while the extracellular matrix, macromolecules, and unresolved cellular structures provide an elastic scaffold in the PVS and ECS Ω_*e*_. The semi-permeable cell membrane Γ separates the intracellular and extracellular domains, while allowing for pressure-driven fluid exchange. Within this context and under the assumption of relatively small deformations (58), Biot’s equations of linear poroelasticity predict the evolution in time and distribution in space of the pressure *p* = *p*(***x***, *t*) and displacement vector **d** = **d**(***x***, *t*), for ***x*** ∈ Ω_*i*_, Ω_*e*_ and *t >* 0, and read as:

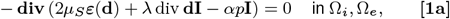

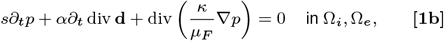

where **div** and div denotes the divergence acting on tensors and vectors, respectively, ***ε***(**d**) is the symmetric strain tensor, **I** is the identity tensor, and ∂_*t*_is the time derivative.

At the cell membrane Γ, a Robin-type interface condition allows for the transmembrane fluid flux 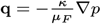 to be regulated by the membrane permeability *L*_*p*_, the hydrostatic pressure difference [*p*] and the osmotic pressure *p*_osm_:

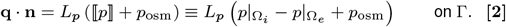

For the scenario with an osmotic driving force, we link the osmotic pressure with the concentration difference Δ*c* by the van’t Hoff equation *p*_osm_ = Δ*ciRT*, where *i* is the van’t Hoff index (*i* = 1 for glucose), *R* is the ideal gas constant and *T* denotes the absolute temperature. For all other scenarios we set *p*_osm_ = 0. Further, imposing mass conservation, continuity of the displacement and momentum conservation on the cell membrane yields the additional constraints

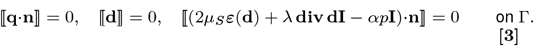

These equations are parameterized by the Lamé constants *µ*_*S*_ *>* 0 and *λ >* 0, the Biot-Willis coefficient *α* ∈ (0, 1], the storage coefficient *s* ≥ 0, the permeability *κ >* 0, the fluid viscosity *µ*_*F*_ ≥ 0 and the hydraulic conductivity of the cell membrane *L*_*p*_. Material parameter values were drawn from experimental literature where available (Table S1). For parameters lacking specific in vivo measurements, such as the permeability of the endfoot cytoskeleton, values were estimated to be consistent with adjacent tissue properties. In particular, we set the Biot-Willis coefficient *α* = 1 as the porous frame is highly compliant compared to its incompressible constituents, and the storage coefficient to *s* = 0 as fluid compressibility is negligible at physiological pressures; a sensitivity analysis (*s* = 10^−9^ Pa^−1^) confirmed this assumption has no impact on the results (Figure S7). Further, we assume that the system starts at rest and at zero pressure.

### Interaction with the surroundings: dynamic boundary conditions

To model the response of the tissue to vascular pulsations, we prescribe a given radial motion of the arteriole wall. Specifically, we assume that the arteriole dilates and constricts rhythmically in time with frequency *f* and amplitude *A* in the radial direction **r** on Γ_*VW*_ (see Table S2), uniformly along the axial direction:

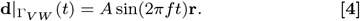

Here, *A* corresponds to a relative diameter change of approximately 0.8 % (*A* = 50 nm) for the cardiac cycle and 8 % (*A* = 500 nm) for vasomotion. We thus neglect any phase shift along the vessel wall, which is a reasonable assumption here considering the high wave speed of the arterial pulse wave relative to the microscopic length of the vessel segment. Moreover, we assume that this interface is impermeable; i.e. we do not allow for fluid flow from the perivascular space into the lumen or vice versa. To also investigate the potential effect of physiological waveform asymmetry, we extracted the cardiac-driven vessel movement data from Mestre et al. (23) (Fig. 3e) in place of Eq. (4). To isolate the effect of the asymmetry from differences in movement amplitude, we scaled the vessel displacement to preserve the same area under the curve as the idealized sinusoidal curve.

At the outer radial boundary of the geometry (Γ_radial_ ≡ Γ_EF−radial_ ∪ Γ _ECS −radial_) we consider a homogenized representation of the neuropil as a soft and permeable tissue with permeability *κ*_far_ and stiffness *E*_far_. We allow for fluid flow across both the outer extracellular and intracellular boundaries. Concretely, we assume a linear pressure profile between the outer boundary and a far-field pressure *p*_far_ at a distance *l*_far_ yielding the condition:

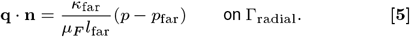

Similarly, we use an elastic spring model for the outer radial displacement: the outer endfoot sheath is allowed to dilate and constrict, but an opposing force linear to the displacement magnitude mimics the resistance of the surrounding tissue.

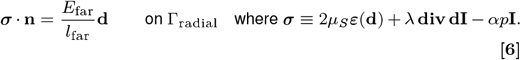

We fix the axial boundaries of the intracellular and non-perivascular extracellular spaces by not allowing for any displacement in the axial direction or any fluid flow there. More precisely, we set

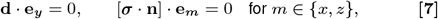

with the standard basis vectors **e**_*x*_, **e**_*y*_and **e**_*z*_, and noting that the vessel centerline is oriented in the y-direction. However, for the axial perivascular boundary, we consider two scenarios. One side remains closed, but we set the other side to be either open (by prescribing a zero pressure) or closed (by prescribing zero flow). These two scenarios model the two extreme cases of either an infinitely small or an infinitely large resistance of the PVS fluid pathway. For a summary of all boundary conditions, see Table S2.

### Numerical approximation and simulation software

To solve the Biot equations Eq. (1) numerically, we employ a three-field formulation with the fluid flux 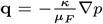 as additional unknown (59, 60). Discretizing in time with a finite-difference approach, we use a non-conforming discontinuous Galerkin discretization in space, which ensures local (fluid) mass conservation. This numerical approximation property is especially relevant here, as the fluid mass and the associated changes in pressure and volume are primary quantities of interest. Moreover, the scheme provides optimal (first-order) convergence rates at a low number of degrees of freedom per element (60), and is thus well-suited for high-resolution meshes of complex geometries. The total number of degrees of freedom is 12 954 440 at each time step. A comprehensive description of the numerical method can be found in the Supplementary Information. We implemented the numerical solution algorithms using the FEniCS finite element software (61), and the complete simulation pipeline is openly available (62).

## Supporting information

displacement movie

pressure movie

## ACKNOWLEDGMENTS

M.E.R. acknowledges support from Stiftelsen Kristian Gerhard Jebsen via the K. G. Jebsen Centre for Brain Fluid Research and Wellcome via Award 313298/Z/24/Z. M.C. and M.E.R. acknowledge support from the Research Council of Norway (RCN) via grant #324239 (EMIx). M.C. acknowledges support from the European Research Council via grant #101141807 (aCleanBrain). R.E. received funding from the Chan Zuckerberg Initiative DAF, an advised fund of Silicon Valley Community Foundation (grand number 2023-331840), the South-Eastern Norway Regional Health Authority (grant #2025018 and #2024079) and the Letten Foundation. Simulations were performed on the Experimental Infrastructure for Exploration of Exascale Computing (eX3), which is financially supported by the RCN under contract 270053. Figures 1B, 3A, 4A, S1A, S4A were created in BioRender.

## Notes

### Competing Interest Statement

The authors have declared no competing interest.

### Summary of Updates

shorten and fit to PNAS template, add supplementary

