## supplementary material for "Stretch and flow at the gliovascular interface: high-fidelity modelling of astrocyte endfeet"

### Supplementary information for Stretch and flow at the gliovascular interface: high-fidelity modelling of astrocyte endfeet

**Marius Causemann[1], Rune Enger[2,3,4], and Marie E. Rognes[1,4,*]**

[1]Department of Numerical Analysis and Scientific Computing, Simula Research Laboratory, Kristian Augusts gate 23, 0164 Oslo, Norway
[2]Institute of Basic Medical Sciences, University of Oslo, Oslo, Norway
[3]Department of Neurosurgery, Oslo University Hospital - Rikshospitalet, Oslo, Norway
[4]K. G. Jebsen Centre for Brain Fluid Research, Oslo, Norway
[*]

#### ABSTRACT

This document includes supplementary methods, tables, figures, videos, and results accompanying the main text of *Stretch and flow at the gliovascular interface: high-fidelity modelling of astrocyte endfeet*.

#### S1 Supplementary videos

This section includes videos SV1 and SV2.

**Video SV1:** Dynamic displacement induced by wall pulsations

**Video SV2:** Dynamic pressure induced by wall pulsations

#### S2 Supplementary results and figures

This section includes supplementary results, extended descriptions of results presented in the main text, and supplementary figures. Sections presenting extended description or a single supplementary figure are labeled as in the main text for easy retrieval of information.

##### S2.1 Robustness and sensitivity to perivascular pathway resistance and waveforms

This supplementary section includes Figure S1.

##### S2.2 Asymmetric dilation waveforms have a negligible effect on gliovascular coupling

To investigate the potential effect of physiological waveform asymmetry, we extracted the cardiac-driven vessel movement data from Mestre et al. [1](Fig. 3e). To isolate the effect of the asymmetry from differences in movement amplitude, we scaled the vessel displacement to preserve either the same area under the curve (AUC), or the same maximum dilation (amplitude scaled) compared to the idealized sinusoidal curve (Figure S2). Comparing results from these three vessel dilations as drivers of motion reveals that stresses, pressure differences and mean displacements in the different model compartments are not affected by waveform asymmetry, if the maximum amplitude is kept constant (Figure S3). In case of the asymmetric, AUC-scaled curve, we find an increase in these quantities, which is proportional to the increase in absolute vessel displacement magnitude of the AUC-scaled curve. In terms of flow rates and velocities, differences are slightly larger, with peak PVS flow velocity in the asymmetric case reaching up to twice as high as the symmetric case. However, the resulting flow rates differences between all three models stay below 20 %. This behavior reflects the steeper, but shorter vessel expansion in the asymmetric waveforms. We conclude that while certain differences in peak flow quantities exist, the overall behavior of idealized symmetric and asymmetric waveforms is largely equivalent, if differences in magnitude are accounted for.

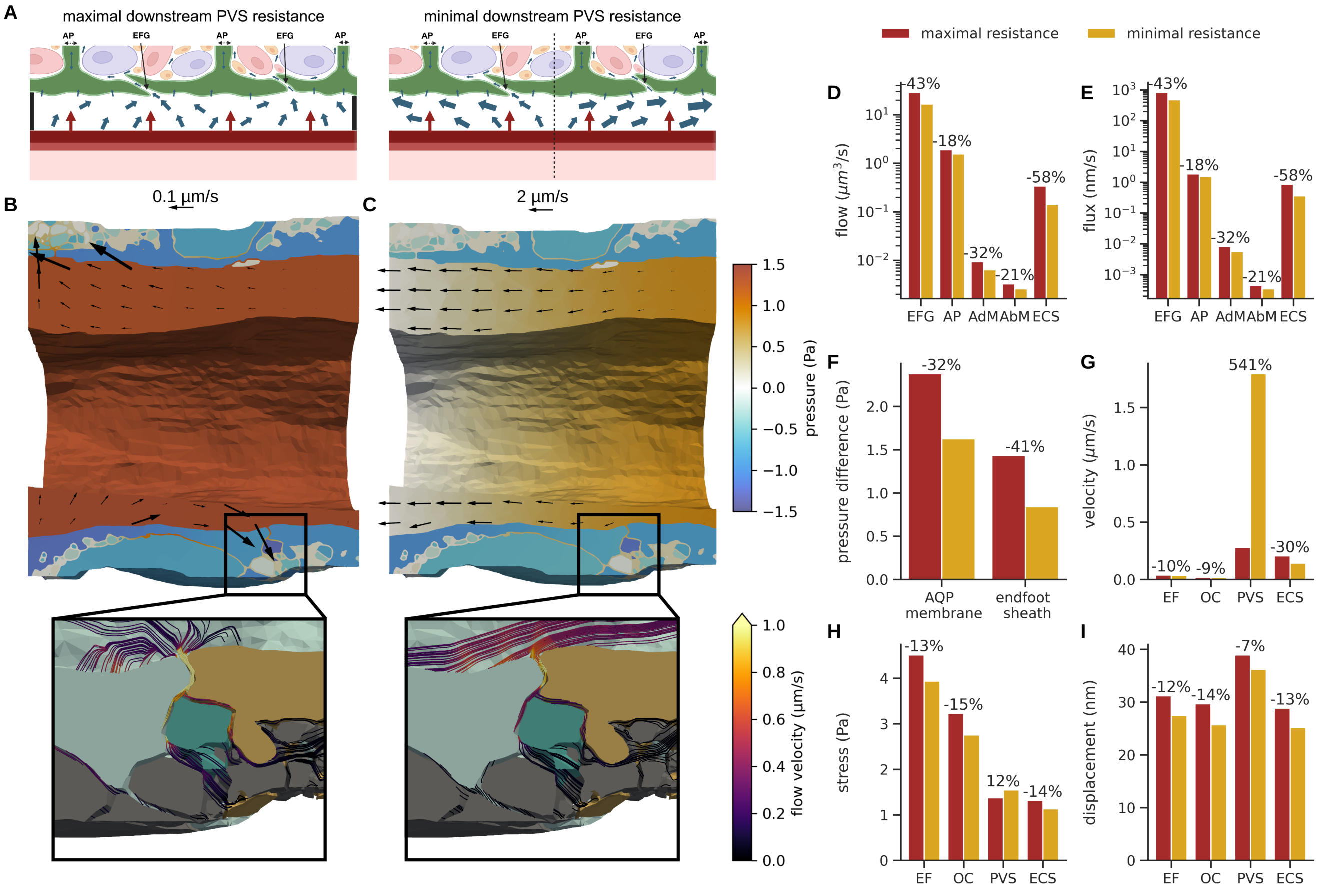


**Figure S1. The downstream PVS resistance affects local flow patterns and tissue motion.** A) Conceptual illustration of minimal and maximal downstream PVS resistance, B–C) fluid pressure and velocity at peak systole for the maximal downstream PVS resistance (B) and minimal PVS flow resistance scenarios (C), with zoom-ins showing streamline visualizations of extracellular fluid flow; D) peak fluid flow rates across the endfoot gaps (EFG, interface between PVS and ECS), the outer astrocyte processes (AP, outer radial boundary of the endfeet), the adluminal endfoot membrane (AdM), the abluminal endfoot membrane (AbM), and the ECS outer radial boundary (ECS); E) peak flux rates (labels as in D); F) peak mean pressure gradient across the adluminal endfoot membrane and the whole endfoot sheath; G) peak mean flow velocities in each compartment; H) peak mean von Mises stress in each compartment; I) peak mean displacement magnitude in each compartment.

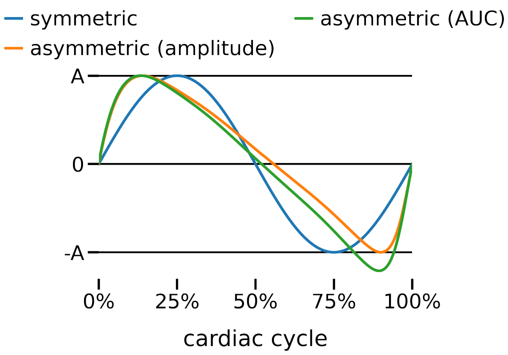


**Figure S2. Comparison of dilation waveforms.** The symmetric waveform is the sine-curve used throughout the paper, while the asymmetric was extracted from Mestre et al. [1](Fig. 3e), and scaled to either match the amplitude or the area under the curve (AUC) of the symmetric waveform.

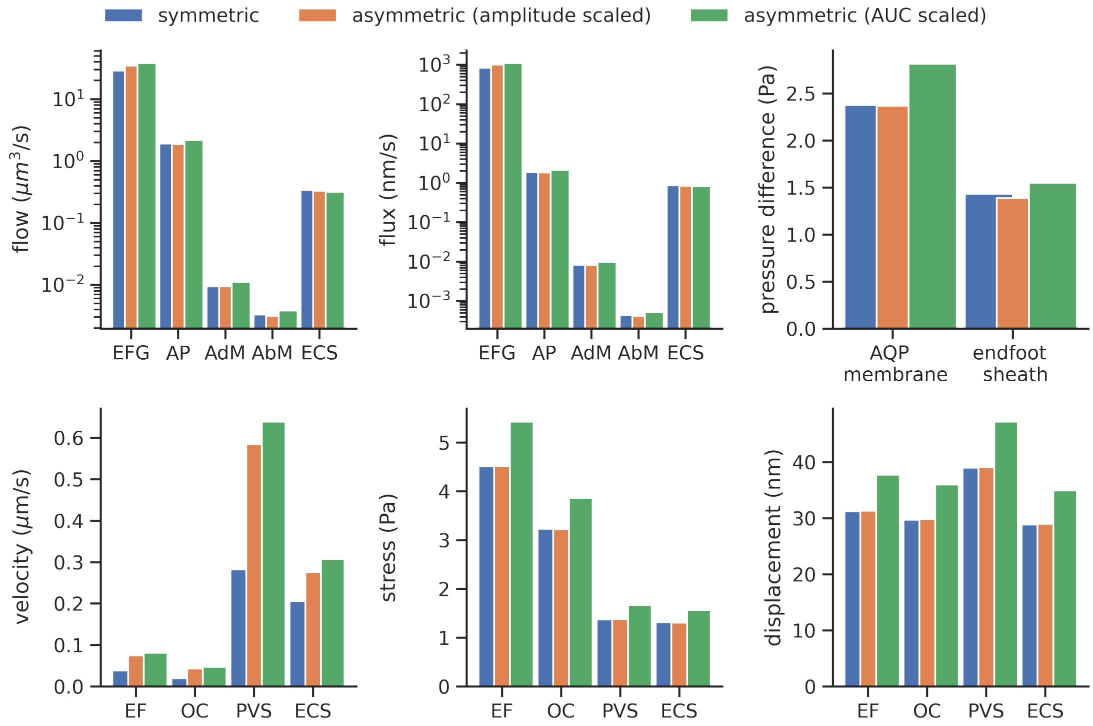


**Figure S3. Comparison of results with symmetric and asymmetric dilation waveforms.** Peak fluid flow rates across the endfoot gaps (EFG, interface between PVS and ECS), the outer astrocyte process (AP, outer radial boundary of the endfeet), the adluminal endfoot membrane (AdM), the abluminal endfoot membrane (AbM) and the ECS outer radial boundary (ECS); peak flux rates; peak mean pressure difference across the adluminal endfoot membrane and the whole endfoot sheath; peak mean flow velocities in each compartment; peak mean von Mises stress in each compartment; peak mean displacement magnitude in each compartment;

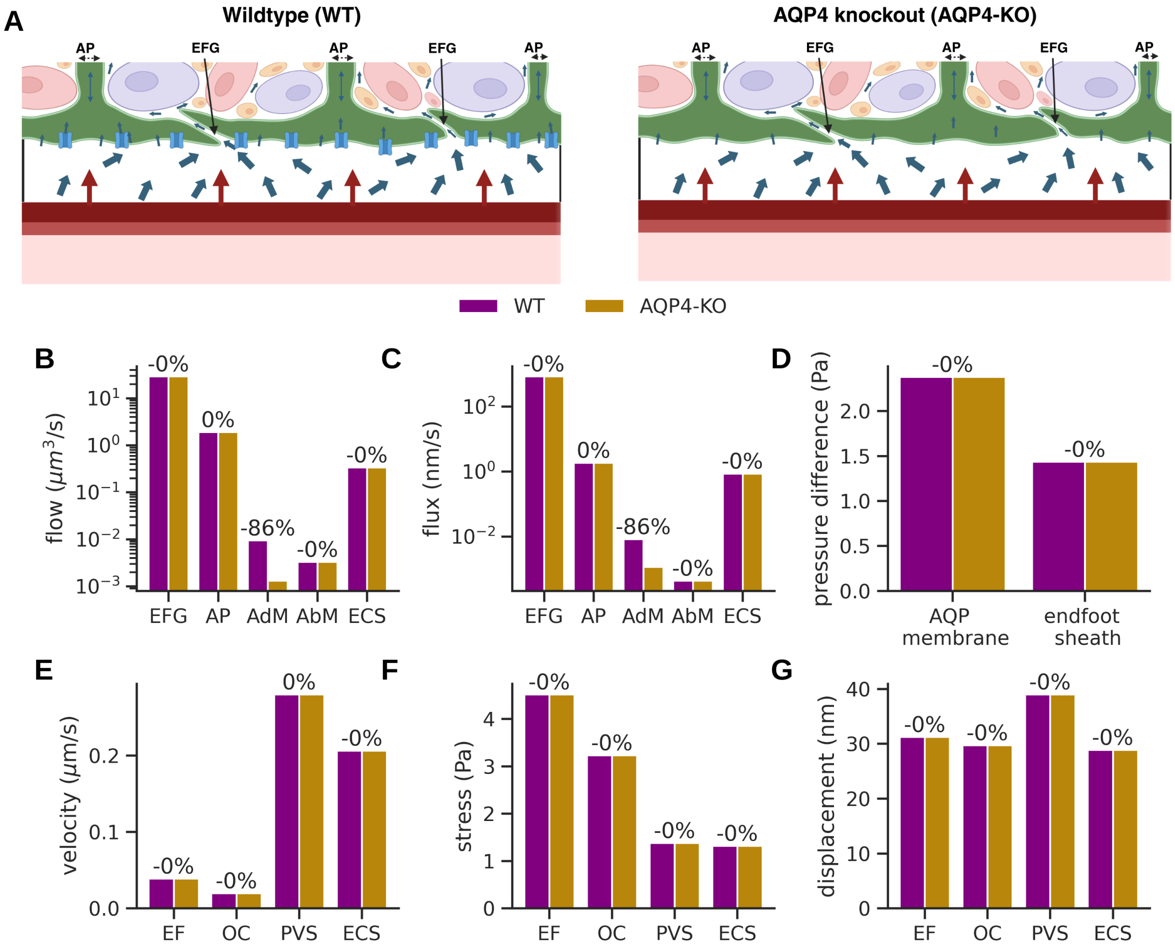


**Figure S4. AQP4-mediated increase in water channel permeability has a negligible impact on gliovascular dynamics.** A) Conceptual illustration of the computational wildtype and AQP4 knockout models; B) peak fluid flow rates across the endfoot gaps (EFG, interface between PVS and ECS), the outer astrocyte process (AP, outer radial boundary of the endfeet), the adluminal endfoot membrane (AdM), the abluminal endfoot membrane (AbM) and the ECS outer radial boundary (ECS); C) peak flux rates (labels as in B); D) peak mean pressure gradient across the adluminal endfoot membrane and the whole endfoot sheath; E) peak mean flow velocities in each compartment; F) peak mean von Mises stress in each compartment; G) peak mean displacement magnitude in each compartment;

##### S2.3 Astrocytic endfoot response to vascular pulsations is resilient to changes in endfoot water permeability

This section includes Figure S4.

##### S2.4 Low-frequency, high-amplitude vasomotion drives large tissue displacement and fluid exchange

To investigate the effects of neurovascular coupling on gliovascular mechanics, we simulated a "vasomotion" scenario characterized by low-frequency (0.2 Hz), high-amplitude (8% dilation) vessel wall motion, mimicking the functional hyperemia observed in response to whisker stimulation [2] or sleep-state dependent vasomotion [3]. We compared this to our baseline "cardiac" scenario (10 Hz, 0.8% dilation) to quantify the impact of high-amplitude vasomotion relative to baseline cardiac pulsations. Consistent with the large (10-fold) increase in arterial dilation amplitude, our model predicts a substantial increase in the mechanical response of the surrounding tissue. Peak displacement magnitudes increased by 877% in the endfeet and 900% in the ECS compared to the cardiac baseline (Figure S5), reaching up to approximately 300–400 nm. The mechanical stress exerted on the gliovascular interface increased drastically, with von Mises stress rising by 827% in the endfeet and 934% in the PVS.

Counter-intuitively, this large increase in driving amplitude did not uniformly amplify fluid velocities. While fluid efflux from the ECS into the surrounding tissue increased substantially (3288%), the flow through the inter-endfoot gaps (EFG) actually decreased by 8% (Figure S5A-B). Similarly, the mean fluid velocity within the ECS increased by only 50%. Visual inspection of the flow fields confirms this observation: velocity magnitude maps reveal that the internal flow patterns and magnitudes remain remarkably similar between the two regimes (Figure S5C), indicating that the 10-fold increase in wall motion does not translate into a proportional increase in local fluid velocity. The disparity is explained by the frequency-dependent poroelastic response of the tissue. In the high-frequency cardiac regime (10 Hz), the rapid wall motion prevents the ECS pressure from equilibrating with that of the PVS, creating a phase lag between the compartments. This lag generates a transient pressure gradient that drives fluid through the endfoot gaps (Figure S5B). Conversely, the slower dilation of the vasomotion regime (0.2 Hz) allows sufficient time for pressure equilibration. Consequently, the PVS and ECS pressures become almost synchronized, preventing the formation of large gradients across the endfoot layer and limiting the amplification of local flow velocities (Figure S5B).

However, while the spatial gradients remain comparable, the absolute pressure amplitude rises substantially during vasomotion. This global pressurization drives increased exchange across high-resistance pathways, which are governed by the pressure difference relative to the far-field or the intracellular pressure. Transmembrane fluxes (AdM and AbM) thus increased by 696–1453%, and most prominently, the fluid efflux into the surrounding tissue increased by 3288%, due to the combined effect of larger absolute pressure changes, and pressure synchronization between PVS and ECS (Figure S5A). The hydraulic impact of vasomotion becomes even more evident when considering the total displaced fluid volume per cycle. Due to the lower frequency, the duration of the expansion phase increases 50-fold (2.5 s vs 0.05 s). Consequently, even with comparable flow rates, the integrated volume of fluid transported across the interface increases substantially. The flow volume passing through the endfoot gaps increased 42-fold, while the volume exchanged across the cell membrane and with the surrounding ECS increased by 300–753-fold (Figure S5A, middle). Thus, vasomotion displaces much higher fluid volumes, primarily by sustaining flow for longer durations, rather than by amplifying peak velocities.

##### S2.5 Astrocytic endfoot response to vasomotion is resilient to changes in endfoot water permeability

To confirm our observation of minor effects of reduced endfoot water permeability (AQP4-knockout model) on gliovascular dynamics from the cardiac setting, we performed additional simulations with vasomotion frequency (0.2 Hz) and amplitude (8% dilation) and approximately sevenfold decreased permeability of the adluminal endfoot membrane. Compared with the baseline vasomotion case, flow across the adluminal endfoot membrane decreased by a factor of seven, but did not induce any substantial changes in tissue motion or other flow rates (all below 1% relative difference) due to the small magnitude of the transmembrane flux (Figure S6).

##### S2.6 Sensitivity to the storage coefficient $s$

In our primary analysis, we set the storage coefficient to $s = 0$, assuming that the fluid and solid constituents are intrinsically incompressible. To examine this assumption, we performed an additional numerical experiment using $s = 10^{-9}\,\mathrm{Pa}^{-1}$. This value was chosen as a conservative upper bound for the effective compressibility of the fluid-saturated tissue. It accounts for the compressibility of pure water ($\approx 4.6 \times 10^{-10}\,\mathrm{Pa}^{-1}$) as well as the proteins constituting the extracellular matrix and cytoskeleton. We found that introducing this compressibility term yielded no discernible difference in pressure, displacement, or flow predictions compared to the baseline ($s = 0$) model (Figure S7).

#### S3 Supplementary methods

These sections provide a complete description of the mathematical model and numerical approximations employed. All material parameters are given in Table S1 and a summary of boundary conditions in Table S2.

##### S3.1 Three-field formulation of the cell-by-cell dual poroelasticity model

We restate the system of coupled poroelasticity equations with the fluid flux $\mathbf{q} = -\frac{\kappa}{\mu_F}\nabla p$ as an additional unknown. The equations then read: find the displacement $\mathbf{d} : \Omega_i \cup \Omega_e \to \mathbb{R}$, the fluid pressure $p : \Omega_i \cup \Omega_e \to \mathbb{R}$, and the fluid flux $\mathbf{q} : \Omega_i \cup \Omega_e \to \mathbb{R}$ such that

$$
\begin{aligned}
-\mathbf{div}\,(2\mu_S \boldsymbol{\varepsilon}(\mathbf{d}) + \lambda\,\mathrm{div}\,\mathbf{d}\mathbf{I} - \alpha p \mathbf{I}) &= 0, \\
\mathbf{q} + \frac{\kappa}{\mu_F}\nabla p &= 0, \\
s\partial_t p + \alpha \partial_t\,\mathrm{div}\,\mathbf{d} + \mathrm{div}\,\mathbf{q} &= 0,
\end{aligned}
\tag{1}
$$

all over $\Omega_i, \Omega_e$. The interface conditions on the cell membrane $\Gamma = \overline{\Omega_i} \cap \overline{\Omega_e}$ read as

$$
\mathbf{q}\cdot\mathbf{n} = L_p\,(\llbracket p \rrbracket + p_{\mathrm{osm}}), \quad \llbracket \mathbf{q}\cdot\mathbf{n} \rrbracket = 0, \quad \llbracket \mathbf{d} \rrbracket = 0, \quad \llbracket (2\mu_S \boldsymbol{\varepsilon}(\mathbf{d}) + \lambda\,\mathrm{div}\,\mathbf{d}\mathbf{I} - \alpha p \mathbf{I})\cdot\mathbf{n} \rrbracket = 0, \tag{2}
$$

where the jump $\llbracket \cdot \rrbracket$ denotes the difference over $\Gamma$. The complete set of dynamic boundary conditions are as follows, for $t > 0$.

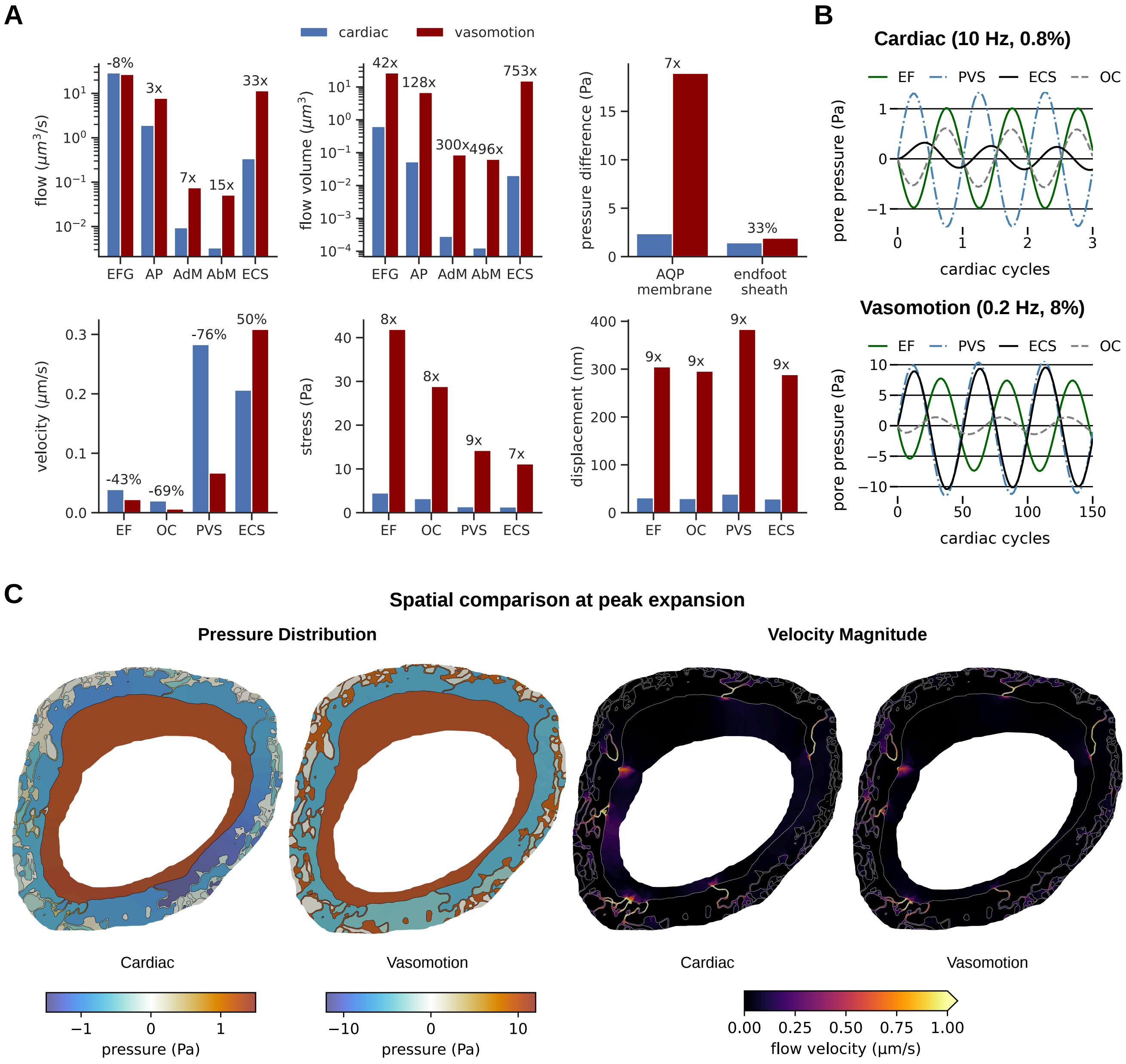


**Figure S5.** **Vasomotion drives large tissue displacement, but does not uniformly amplify local fluid dynamics. (A)** Relative change in key mechanical and fluid dynamical quantities during vasomotion (0.2 Hz, 8% dilation) compared to the cardiac baseline (10 Hz, 0.8% dilation). **(B)** Temporal traces of pore pressure in the PVS, ECS, and endfeet (EF). **(C)** Cross-sectional maps of pore pressure and fluid velocity magnitude at peak expansion.

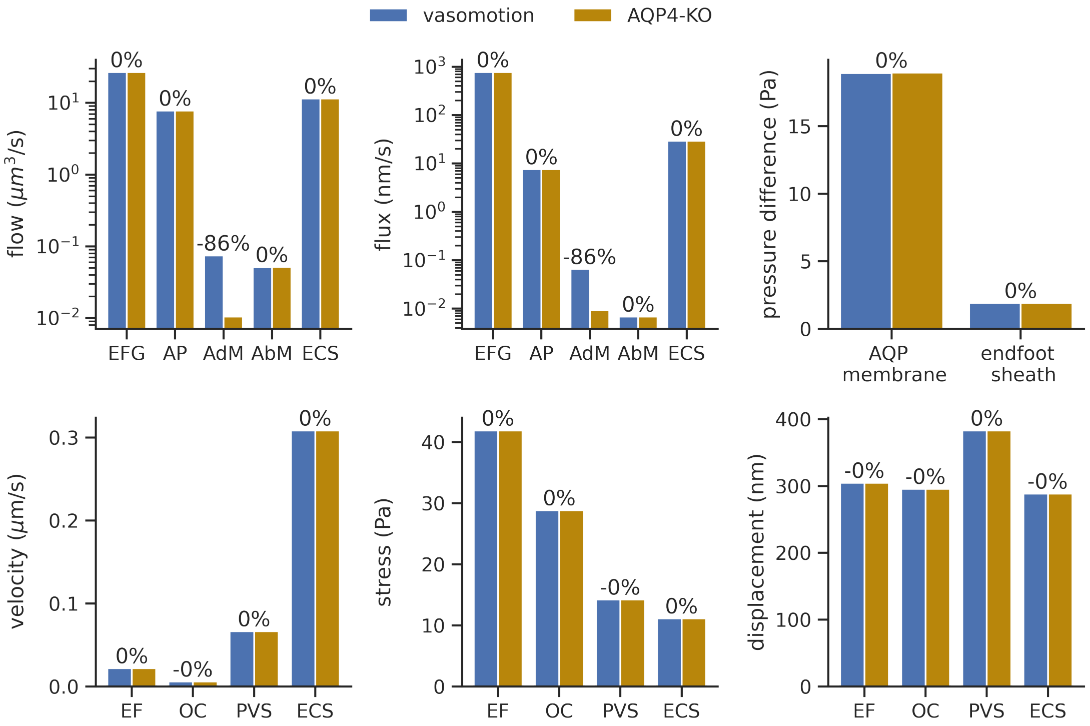


**Figure S6. Comparison of vasomotion results for wildtype and AQP4-knockout models.** Peak fluid flow rates across the endfoot gaps (EFG, interface between PVS and ECS), the outer astrocyte process (AP, outer radial boundary of the endfeet), the adluminal endfoot membrane (AdM), the abluminal endfoot membrane (AbM) and the ECS outer radial boundary (ECS); peak flux rates; peak mean pressure difference across the adluminal endfoot membrane and the whole endfoot sheath; peak mean flow velocities in each compartment; peak mean von Mises stress in each compartment; peak mean displacement magnitude in each compartment;

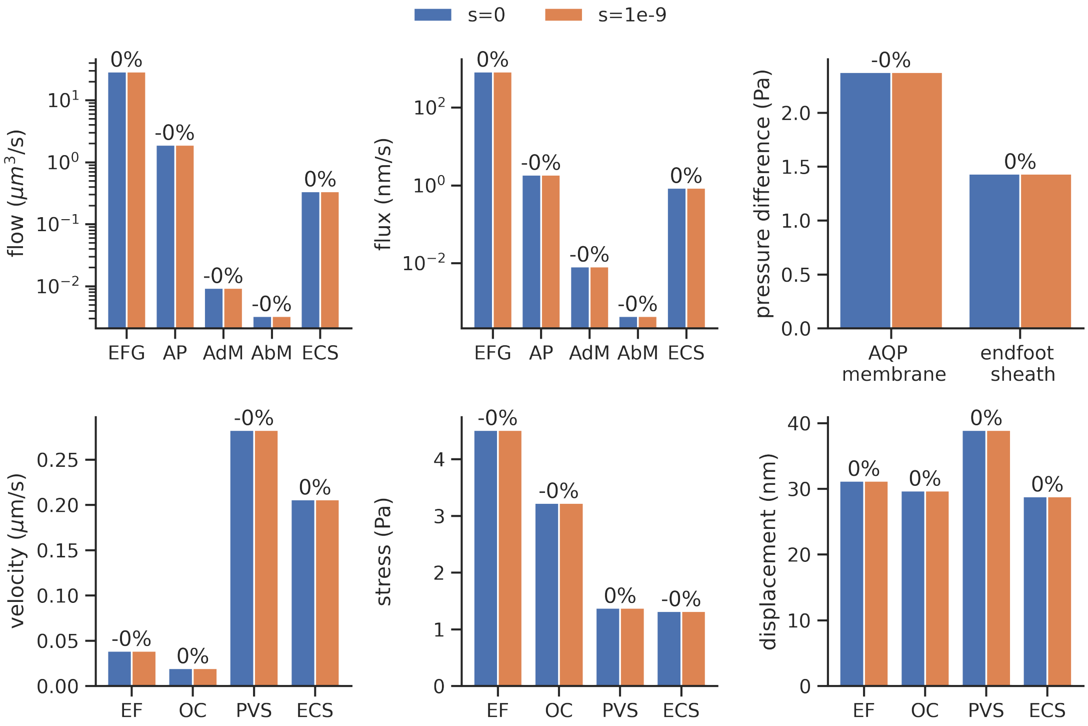


**Figure S7. Comparison of results with storage coefficient $s = 0$ and $s = 10^{-9}$ Pa$^{-1}$.** Peak fluid flow rates across the endfoot gaps (EFG, interface between PVS and ECS), the outer astrocyte process (AP, outer radial boundary of the endfeet), the adluminal endfoot membrane (AdM), the abluminal endfoot membrane (AbM) and the ECS outer radial boundary (ECS); peak flux rates; peak mean pressure difference across the adluminal endfoot membrane and the whole endfoot sheath; peak mean flow velocities in each compartment; peak mean von Mises stress in each compartment; peak mean displacement magnitude in each compartment;

| Parameter | Value | Literature values | References / Basis |
|---|---|---|---|
| Stiffness (Young modulus) $E$ | | | |
| – endfeet $E_{\mathrm{EF}}$ | 1500 Pa | 200 – 2650 Pa | [4, 5] |
| – ECS $E_{\mathrm{ECS}}$ | 150 Pa | – | assumed equal to $E_{\mathrm{PVS}}$ |
| – PVS $E_{\mathrm{PVS}}$ | 150 Pa | 60 Pa | [6] |
| Poisson's ratio $\nu$ | 0.3 | 0.3 | [7, 8] |
| Fluid viscosity $\mu_F$ | $0.0007\,\mathrm{Pa\cdot s}$ | $0.0007\,\mathrm{Pa\cdot s}$ | [9] |
| Biot–Willis coefficient $\alpha$ | 1 | 1 | [7] |
| Storage coefficient $s$ | $0\,\mathrm{Pa}^{-1}$ | $0\,\mathrm{Pa}^{-1}$ | [7] |
| Permeability $\kappa$ | | | |
| – endfeet $\kappa_{\mathrm{EF}}$ | $1\cdot 10^{-15}\,\mathrm{m}^2$ | – | scaled relative to $\kappa_{\mathrm{ECS}}$ ($\kappa_{\mathrm{ECS}}\times 0.5$) |
| – ECS $\kappa_{\mathrm{ECS}}$ | $2\cdot 10^{-15}\,\mathrm{m}^2$ | $2\cdot 10^{-15}\,\mathrm{m}^2$ | [6] |
| – PVS $\kappa_{\mathrm{PVS}}$ | $2\cdot 10^{-14}\,\mathrm{m}^2$ | $2\cdot 10^{-14} - 1.8\cdot 10^{-15}\,\mathrm{m}^2$ | [6, 10] |
| Membrane conductance $L_p$ | | | |
| – AQP | $3.49\cdot 10^{-12}\,\mathrm{m\,s^{-1}\,Pa^{-1}}$ | $3.49\cdot 10^{-12}\,\mathrm{m\,s^{-1}\,Pa^{-1}}$ | [11] |
| – non-AQP | $4.88\cdot 10^{-13}\,\mathrm{m\,s^{-1}\,Pa^{-1}}$ | $4.88\cdot 10^{-13}\,\mathrm{m\,s^{-1}\,Pa^{-1}}$ | [11] |
| Far-field pressure $p_{\mathrm{far}}$ | 0 Pa | - | |
| Far-field distance $l_{\mathrm{far}}$ | 50 µm | - | |
| Outer stiffness $E_{\mathrm{far}}$ | 1500 Pa | $1389 \pm 289$ Pa | [12] |
| Outer permeability $\kappa_{\mathrm{far}}$ | $10^{-16}\,\mathrm{m}^2$ | $10^{-17} - 10^{-15}\,\mathrm{m}^2$ | [13] |
| Stiffer PVS scenario | $E_{\mathrm{PVS}} \times 2, 4, 6, 8$ | - | |

**Table S1.** Material parameters for the in-silico model scenarios. Reported ranges are derived from the cited experimental studies. The Lamé parameters $\mu_S$ and $\lambda$ are related to the Young modulus $E$ and Poisson's ratio $\nu$ by $\lambda = \frac{E\nu}{(1+\nu)(1-2\nu)}$ and $\mu_S = \frac{E}{2(1+\nu)}$.

| Boundary | Label | continuity condition | momentum condition |
|---|---|---|---|
| Interface between lumen and PVS | $\Gamma_{\mathrm{VW}}$ | no flow | prescribed displacement |
| Axial end of the PVS (front) | $\Gamma_{\mathrm{PVS\text{-}A}}$ | no flow | no axial displacement |
| Axial end of the PVS (rear) | $\Gamma_{\mathrm{PVS\text{-}B}}$ | no flow \| zero pressure | no axial displacement |
| Outer radial boundary (ICS) | $\Gamma_{\mathrm{EF\text{-}radial}}$ | far field pressure | linear resistance |
| Outer radial boundary (ECS) | $\Gamma_{\mathrm{ECS\text{-}radial}}$ | far field pressure | linear resistance |
| Axial end of the ECS (front, rear) | $\Gamma_{\mathrm{ECS\text{-}axial}}$ | no flow | no axial displacement |
| Axial end of the ICS (front, rear) | $\Gamma_{\mathrm{EF\text{-}axial}}$ | no flow | no axial displacement |

**Table S2. Summary of model boundaries, boundary conditions and boundary assumptions**. We use the terminology: no flow: $\mathbf{q}\cdot\mathbf{n} = 0$, zero pressure: $p = 0$, and for far-field pressure, prescribed displacement, no axial displacement and linear resistance, see Methods: Interaction with the surroundings: dynamic boundary conditions.

(i) On the inner boundary $\Gamma_{\mathrm{VW}}$, we set:

$$\mathbf{d}(t) = \mathbf{d}_{\mathrm{VW}}(t) \equiv A\sin(2\pi f t)\mathbf{r},$$
$$\mathbf{q}(t)\cdot\mathbf{n} = 0,$$

where $A$ is the amplitude and $f$ the frequency of the vessel wall motion, and $\mathbf{r}$ denotes the unit-length radial direction from the vessel centerline.

(ii) On the axial boundaries of the geometry: $\Gamma_{\mathrm{axial}} \equiv \Gamma_{\mathrm{ECS-axial}} \cup \Gamma_{\mathrm{EF-axial}} \cup \Gamma_{\mathrm{PVS-A}} \cup \Gamma_{\mathrm{PVS-B}}$, we impose no flow:

$$\mathbf{q}(t)\cdot\mathbf{n} = 0, \tag{3}$$

except in the (open) PVS model variation with minimal PVS pathway resistance where we replace (3) on $\Gamma_{\mathrm{PVS-B}}$ by instead imposing $p = 0$.

(iii) Also on $\Gamma_{\mathrm{axial}}$, with the vessel centerline coinciding with the y-direction of the coordinate system, we restrict axial displacement and allow free radial movement by the following conditions

$$\mathbf{d}(t)\cdot\mathbf{e}_y = 0, \qquad \mathbf{e}_m\cdot\boldsymbol{\sigma}(t)\cdot\mathbf{n} = 0 \quad m \in \{x, z\},$$

where the poroelastic stress tensor $\boldsymbol{\sigma} \equiv 2\mu_S\boldsymbol{\varepsilon}(\mathbf{d}) + \lambda\,\mathrm{div}\,\mathbf{d}\mathbf{I} - \alpha p\mathbf{I}$, and $\mathbf{e}_x$, $\mathbf{e}_y$ and $\mathbf{e}_z$ are the unit vectors in the direction of each coordinate axis.

(iv) Assuming no displacement in the far field distance $l_{\mathrm{far}} > 0$, we use a Robin boundary condition on the radial boundaries $\Gamma_{\mathrm{radial}} \equiv \Gamma_{\mathrm{EF-radial}} \cup \Gamma_{\mathrm{ECS-radial}}$, emulating the elastic response of the surrounding tissue:

$$\boldsymbol{\sigma}(t)\cdot\mathbf{n} = \frac{E_{\mathrm{far}}}{l_{\mathrm{far}}}\mathbf{d}(t),$$

where $E_{\mathrm{far}}$ is the elastic Young modulus of the surrounding tissue.

(v) Additionally on $\Gamma_{\mathrm{radial}}$, we allow fluid outflow across both the endfeet and the ECS outer boundaries, assuming a (homogenized) permeability $\kappa_{\mathrm{far}}$ and a far field pressure at a distance $l_{\mathrm{far}}$, resulting in the condition:

$$\mathbf{q}(t)\cdot\mathbf{n} = \frac{\kappa_{\mathrm{far}}}{\mu_F l_{\mathrm{far}}}(p(t) - p_{\mathrm{far}}).$$

##### S3.2 Semi-discrete variational formulation of the three-field cell-by-cell dual poroelasticity model

We introduce the following spaces for the displacements and fluxes, respectively, incorporating the essential boundary conditions (in the sense of traces) as appropriate:

$$\begin{aligned}\mathbf{V}_{\mathbf{g}} &= \{\mathbf{d} \in H^1(\Omega;\mathbb{R}^3) \,:\, \mathbf{d}|_{\Gamma_{\mathrm{VM}}} = \mathbf{g} \text{ and } \mathbf{d}\cdot\mathbf{e}_y|_{\Gamma_{\mathrm{axial}}} = 0\},\\ \mathbf{W} &= \{\mathbf{q} \in H(\mathrm{div},\Omega) \,:\, \mathbf{q}\cdot\mathbf{n}|_{\Gamma_{\mathrm{VW}}\cup\Gamma_{\mathrm{axial}}} = 0\}.\end{aligned} \tag{4}$$

In (4), $H^1(\Omega;\mathbb{R}^3)$ and $H(\mathrm{div},\Omega)$ are the standard Sobolov spaces of square-integrable vector-valued functions with square-integrable gradient components and divergence, respectively. We denote the $L^2(\Omega)$-inner product as

$$(a,b)_\Omega \equiv \int_\Omega a\cdot b\,\mathrm{d}\boldsymbol{x} \tag{5}$$

and omit the subscript $\Omega$ when $\Omega = \Omega_i \cup \Omega_e$.

After time discretization with an implicit Euler method, the time-dependent problem, equipped with the initial conditions

$$\mathbf{d}^0 = \mathbf{0} \quad \text{and} \quad p^0 = 0,$$

simplifies to the following sequence of stationary, semi-discrete variational problems: for $n = 1, 2, 3, \ldots$, with time step size $\tau^n \equiv t^n - t^{n-1} > 0$, find $\mathbf{d}^n \in \mathbf{V}_{\mathbf{d}_{\mathrm{VW}}(t^n)}$, $\mathbf{q}^n \in \mathbf{W}$ and $p^n \in L^2(\Omega)$ satisfying

$$(2\mu_S\boldsymbol{\varepsilon}(\mathbf{d}^n), \boldsymbol{\varepsilon}(\mathbf{v})) + \lambda(\mathrm{div}\,\mathbf{d}^n, \mathrm{div}\,\mathbf{v}) - (\alpha p^n, \mathrm{div}\,\mathbf{v}) + r_d(\mathbf{d}^n, \mathbf{v}) = 0 \qquad \forall \mathbf{v} \in \mathbf{V_0} \tag{6a}$$

$$(K^{-1}\mathbf{q}^n, \mathbf{w}) - (p^n, \mathrm{div}\,\mathbf{w}) + (L_p^{-1}\mathbf{q}^n\cdot\mathbf{n}, \mathbf{w}\cdot\mathbf{n})_\Gamma + r_q(\mathbf{q}^n, \mathbf{w}) = (p_{\mathrm{osm}}, \mathbf{w}\cdot\mathbf{n})_\Gamma \qquad \forall \mathbf{w} \in \mathbf{W} \tag{6b}$$

$$(\alpha\tau^{-1}\,\mathrm{div}\,\mathbf{d}^n, \phi) + (\mathrm{div}\,\mathbf{q}^n, \phi) + (s\tau^{-1}p^n, \phi) = (\alpha\tau^{-1}\,\mathrm{div}\,\mathbf{d}^{n-1} + s\tau^{-1}p^{n-1}, \phi) \quad \forall \phi \in L^2(\Omega) \tag{6c}$$

The hydraulic conductivity $K$ is defined by $K = \frac{\kappa}{\mu_F}$. With the far field distance $l_{\text{far}}$, the Young modulus $E_{\text{far}}$ and hydraulic conductivity $K_{\text{far}}$ of the surrounding tissue, the contributions from the Robin boundary conditions are as follows:

$$r_d(\mathbf{d},\mathbf{v}) = (\frac{E_{\text{far}}}{l_{\text{far}}}\mathbf{d},\mathbf{v})_{\Gamma_{\text{radial}}},$$
$$r_q(\mathbf{q},\mathbf{w}) = (K_{\text{far}}^{-1} l_{\text{far}}\mathbf{q}\cdot\mathbf{n},\mathbf{w}\cdot\mathbf{n})_{\Gamma_{\text{radial}}}.$$

Note that the interface term $(L_p^{-1}\mathbf{q}^n\cdot\mathbf{n},\mathbf{w}\cdot\mathbf{n})_\Gamma$ and the Robin term $r_q(\cdot,\cdot)$ in (6b) stems from integration by parts and insertion of the interface flux condition in the following manner:

$$\begin{aligned}
&\int_{\Omega_i\cup\Omega_e} K^{-1}\mathbf{q}^n\cdot\mathbf{w} + \nabla p^n\cdot\mathbf{w}\,\mathrm{d}\boldsymbol{x}\\
&= \int_{\Omega_i\cup\Omega_e} K^{-1}\mathbf{q}^n\cdot\mathbf{w}\,\mathrm{d}\boldsymbol{x} - \int_{\Omega_i\cup\Omega_e} p^n \operatorname{div}\mathbf{w}\,\mathrm{d}\boldsymbol{x} + \int_\Gamma p^n|_{\Omega_i}\mathbf{w}\cdot\mathbf{n}\,\mathrm{d}s - \int_\Gamma p^n|_{\Omega_e}\mathbf{w}\cdot\mathbf{n}\,\mathrm{d}s + \int_{\Gamma_{\text{radial}}} p^n\mathbf{w}\cdot\mathbf{n}\,\mathrm{d}s\\
&= (K^{-1}\mathbf{q}^n,\mathbf{w}) - (p^n,\operatorname{div}\mathbf{w}) + ([\![p^n]\!],\mathbf{w}\cdot\mathbf{n})_\Gamma + (K_{\text{far}}^{-1}l_{\text{far}}\mathbf{q}^n\cdot\mathbf{n},\mathbf{w}\cdot\mathbf{n})_{\Gamma_{\text{radial}}}\\
&\qquad = (K^{-1}\mathbf{q}^n,\mathbf{w}) - (p^n,\operatorname{div}\mathbf{w}) + (L_p^{-1}\mathbf{q}^n\cdot\mathbf{n} - p_{\text{osm}},\mathbf{w}\cdot\mathbf{n})_\Gamma + r(\mathbf{q}^n,\mathbf{w}) \qquad (7)
\end{aligned}$$

for all $\mathbf{w}\in\mathbf{W}$.

##### S3.3 Fully discrete approximation and numerical solution of the cell-by-cell dual poroelasticity model

We consider a mesh $\mathcal{T} = \{E\}$ of $\Omega = \Omega_i\cup\Omega_e$, consisting of tetrahedral mesh cells $E$, and conforming to the domains $\Omega_i$ and $\Omega_e$ and to the cell membrane $\Gamma$. We denote the restriction of $\mathcal{T}$ to $\Omega_i$ and $\Omega_e$ by $\mathcal{T}_i$ and $\mathcal{T}_e$, respectively. The collection of all interior facets (i.e. triangular faces of the tetrahedral mesh cells) in $\mathcal{T}_i$ and $\mathcal{T}_e$ are denoted by $\mathcal{F}_{i,i}$ and $\mathcal{F}_{i,e}$, respectively. We define the union of facets interior both to $\mathcal{T}_i$ and $\mathcal{T}_e$ as $\mathcal{F}_i = \mathcal{F}_{i,i}\cup\mathcal{F}_{i,e}$. Moreover, we denote the exterior facets associated with the boundary parts $\Gamma_{\text{VW}}$, $\Gamma_{\text{PVS-A}}$, $\Gamma_{\text{PVS-B}}$, $\Gamma_{\text{EF-axial}}$, $\Gamma_{\text{EF-radial}}$, $\Gamma_{\text{ECS-axial}}$, $\Gamma_{\text{axial}}$ and $\Gamma_{\text{ECS-radial}}$ with $\mathcal{F}_{\text{VW}}$, $\mathcal{F}_{\text{PVS-A}}$, $\mathcal{F}_{\text{PVS-B}}$, $\mathcal{F}_{\text{EF-axial}}$, $\mathcal{F}_{\text{EF-radial}}$, $\mathcal{F}_{\text{ECS-axial}}$, $\mathcal{F}_{\text{axial}}$ and $\mathcal{F}_{\text{ECS-radial}}$, respectively.

To solve (6) numerically, we utilize a non-conforming discontinuous Galerkin discretization, with a formulation proposed by [14] in the case of a single poroelastic domain. In particular, we use Crouzeix–Raviart elements for the displacements $\mathbf{d}$ (which are only continuous at the facet barycenter and thus are not conforming in $H^1$), the lowest order Raviart–Thomas-Nédélec elements for the Darcy velocity $\mathbf{q}$ and piecewise constants for the fluid pressure $p$, all defined relative to the mesh $\mathcal{T}$. This method is locally mass conservative and locking-free, and its comparably low number of degrees of freedom per mesh cell makes it suitable for high-resolution meshes of complex geometries. More precisely, the non-conforming Crouzeix-Raviart space is defined as [15]:

$$\begin{aligned}
\mathbf{V}_{\mathbf{g},h} = \{\mathbf{d}_h \in L^2(\Omega;\mathbb{R}^3) : \mathbf{d}_h|_E \in \mathcal{P}^1(E)^3 \quad \forall E\in\mathcal{T};\\
\int_e [\![\mathbf{d}_h]\!] = 0 \quad \forall e\in\mathcal{F}_i; \int_e [\![\mathbf{d}_h - \mathbf{g}]\!] = 0 \quad \forall e\in\mathcal{F}_{\text{VW}}; \int_e [\![\mathbf{d}_h\cdot\mathbf{e}_y]\!] = 0 \quad \forall e\in\mathcal{F}_{\text{axial}}\},
\end{aligned} \qquad (8)$$

where $\mathcal{P}^1(E)$ denotes the space of linear polynomials defined over the tetrahedra $E$. Note that the continuity condition $\int_e [\![\mathbf{d}_h]\!] = 0$ for all $e\in\mathcal{F}_i$ is equivalent to continuity at the barycenter of each facet. Further, we set

$$\mathbf{W}_h = \{\mathbf{q}\in H(\operatorname{div},\Omega) : \mathbf{q}|_E \in \mathrm{RT}(E), \quad E\in\mathcal{T}; \quad \mathbf{q}\cdot\mathbf{n}|_{\Gamma_{\text{VW}}\cup\Gamma_{\text{axial}}} = 0\}, \qquad (9)$$

where RT denotes the lowest order Raviart-Thomas-Nédélec elements [16, 17]. Finally, we use the space of piecewise constant functions on each element to approximate the fluid pressure and denote it by $Q_h$.

In order to ensure the coercivity of the bilinear form

$$a(\mathbf{d},\mathbf{v}) = (2\mu_S\boldsymbol{\varepsilon}(\mathbf{d}),\boldsymbol{\varepsilon}(\mathbf{v})) + \lambda(\operatorname{div}\mathbf{d},\operatorname{div}\mathbf{v}), \quad \forall\mathbf{d},\mathbf{v}\in\mathbf{V}_{0,h}, \qquad (10)$$

in the discrete case for the non-conforming Crouzeix-Raviart elements, we include a standard stabilization term [18], which leads to the following modified form:

$$a_h(\mathbf{d},\mathbf{v}) = a(\mathbf{d},\mathbf{v}) + 2\mu_s\eta\sum_{e\in\mathcal{F}_i} h_e^{-1}[\![\mathbf{d}]\!][\![\mathbf{v}]\!] \quad \forall\mathbf{d},\mathbf{v}\in\mathbf{V}_{0,h}. \qquad (11)$$

In (11), $h_e$ denotes the average cell diameter of the cells associated with the edge $e$ and $\eta > 0$ is a penalty parameter. (We choose $\eta = 1$.) As shown by [18] for the equations of linear elasticity, the stabilization ensures positive definiteness of the bilinear form $a_h(\cdot,\cdot)$ and optimal approximation properties of the scheme.

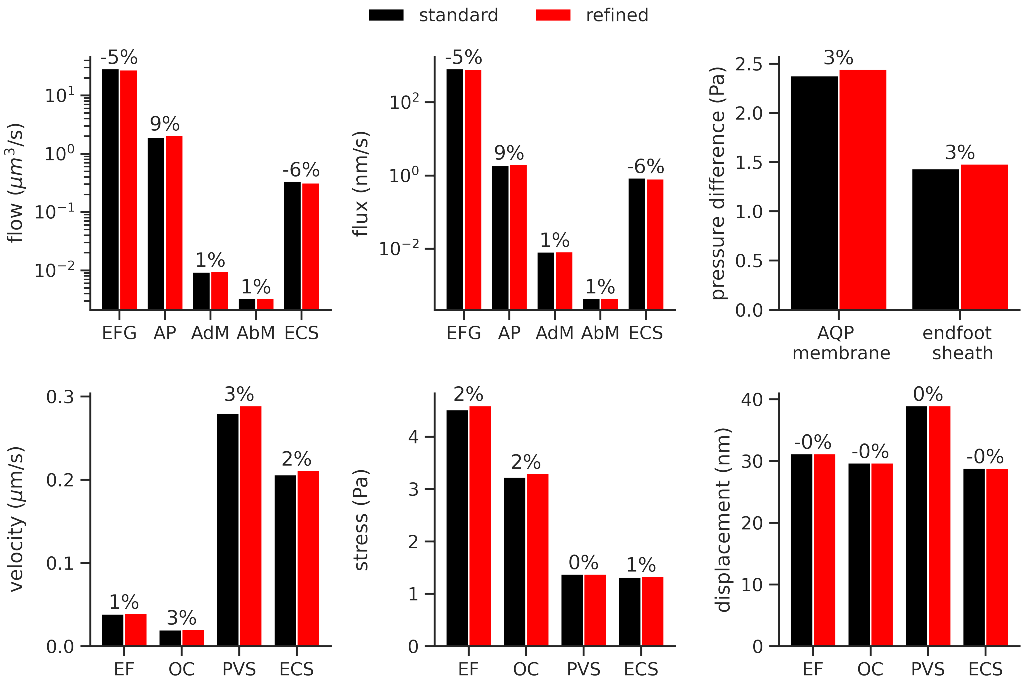


**Figure S8. Verification of computational results.** Numerical verification by comparing key simulated quantities of interest on the standard and a refined mesh with roughly double the number of vertices (258 776 (standard mesh) vs. 494 811 (refined)) and cells (1 425 200 (standard) vs. 2 711 209 (refined)). The percentage above each bar pair (A–F) denotes the relative difference between the quantities computed on the different meshes, revealing only minor differences (9% maximum).

In summary, the fully discrete formulation of (6) with the initial conditions

$$\mathbf{d}_h^0 = \mathbf{d}(0) \quad \text{and} \quad p_h^0 = p(0) \tag{12}$$

then reads: for $n = 1, 2, 3, \dots$, find $\mathbf{d}_h^n \in \mathbf{V}_{\mathbf{d}_{\mathrm{VW}}(t^n),h}$, $\mathbf{q}_h^n \in \mathbf{W}_h$ and $p_h^n \in Q_h$, such that:

$$\begin{aligned} a_h(\mathbf{d}_h^n, \mathbf{v}) - (\alpha p_h^n, \operatorname{div} \mathbf{v}) + r_d(\mathbf{d}_h^n, \mathbf{v}) &= 0 && \forall \mathbf{v} \in \mathbf{V}_{0,h} \\ (K^{-1}\mathbf{q}_h^n, \mathbf{w}) - (p_h^n, \operatorname{div} \mathbf{w}) + (L_p^{-1}\mathbf{q}_h^n \cdot \mathbf{n}, \mathbf{w} \cdot \mathbf{n})_\Gamma + r_q(\mathbf{q}_h^n, \mathbf{w}) &= (p_{\mathrm{osm}}, \mathbf{w} \cdot \mathbf{n})_\Gamma && \forall \mathbf{w} \in \mathbf{W}_h \\ (\alpha\tau^{-1} \operatorname{div} \mathbf{d}_h^n, \phi) + (\operatorname{div} \mathbf{q}_h^n, \phi) + (s\tau^{-1} p_h^n, \phi) &= (\alpha\tau^{-1} \operatorname{div} \mathbf{d}_h^{n-1} + s\tau^{-1} p_h^{n-1}, \phi) && \forall \phi \in Q_h. \end{aligned} \tag{13}$$

Stability and optimal order a-priori error estimates of this scheme have been shown by [14] for the single domain case. Our numerical results, in part included in the Numerical verification section below, indicate that such also hold for the coupled poroelastic problem considered here.

##### S3.4 Numerical verification

We assess the numerical accuracy and convergence of our simulation results by performing experiments with different spatial resolutions. Specifically, we generate a second mesh with roughly twice the number of computational mesh vertices (standard mesh: 158,239 vs. refined: 333,034) and cells (standard: 798,272 vs. refined: 1,704,408), solve the baseline model on both meshes, and compare the results across meshes for a set of key quantities of interest (Figure S8). Finding only minor differences between the two meshes (all relative differences below 10 %, most below 5 %), we conclude that the standard resolution mesh offers sufficient accuracy for our simulations.
